# LYMTACs: Chimeric Small Molecules Repurpose Lysosomal Membrane Proteins for Target Protein Relocalization and Degradation

**DOI:** 10.1101/2024.09.08.611923

**Authors:** Dhanusha A. Nalawansha, Georgios Mazis, Gitte Husemoen, Kate S. Ashton, Weixian Deng, Ryan P. Wurz, Anh T. Tran, Brian A. Lanman, Jiansong Xie, Robert G. Guenette, Shiqian Li, Christopher E. Smith, Suresh Archunan, Manoj K. Agnihotram, Arghya Sadhukhan, Rajiv Kapoor, Sajjan Koirala, Felipe De Sousa E Melo, Patrick Ryan Potts

## Abstract

Proximity-inducing modalities that co-opt cellular pathways offer new opportunities to regulate oncogenic drivers. Inspired by the success of proximity-based chimeras in both intracellular and extracellular target space, here we describe the development of <u>LY</u>sosome <u>M</u>embrane <u>TA</u>rgeting <u>C</u>himera<u>s</u> (LYMTACs) as a novel small molecule-based platform that functions intracellularly to modulate the membrane proteome. Conceptually, LYMTACs are heterobifunctional small molecules that co-opt short-lived lysosomal membrane proteins (LMPs) as effectors to deliver targets for lysosomal degradation. We demonstrate that a promiscuous kinase inhibitor-based LYMTAC selectively targets membrane proteins for lysosomal degradation via RNF152, a short-lived LMP. To extend these findings, we show that oncogenic, membrane-associated KRAS^G12D^ protein can be tethered to RNF152, inducing KRAS relocalization to the lysosomal membrane, inhibiting downstream phospho-ERK signaling, and leading to lysosomal degradation of KRAS^G12D^ in a LYMTAC-dependent manner. Notably, potent cell killing could be attributed to the multi-pharmacology displayed by LYMTACs, which differentiates the LYMTAC technology from existing modalities. Thus, LYMTACs represent a proximity-based therapeutic approach that promises to expand the target space for challenging membrane proteins through targeted protein relocalization and degradation.

## Introduction

Plasma membrane-bound proteins play a central role in cellular signaling, and their deregulation has been implicated in a wide variety of diseases.^1–2^ While the majority of FDA approved drugs engage the membrane proteome, a considerable proportion of membrane-bound targets remain undruggable by current targeted approaches, which are largely comprised of antibodies and small molecule inhibitors (SMI).^3–5^ Despite the potential clinical benefit in modulating membrane proteins, small molecule and antibody-based therapies often suffer from limitations such as stoichiometric target inactivation, challenges in target engagement due to lack of druggable sites, dose-dependent toxicities, and resistance mechanisms.^6–7^ Hence, alternative therapeutic strategies are needed to expand the addressable pool of membrane targets more effectively. Of particular interest, targeted protein degradation (TPD) by proteolysis targeting chimeras (PROTACs) has demonstrated several advantages over inhibitor-based therapies.^8–14^ For instance, PROTACs can bind anywhere on the protein of interest, as opposed to inhibitors, which typically need to bind to an active or allosteric site. Second, degradation of a target removes the entire protein, including its scaffolding functions, essentially mimicking a genetic knockout. Even though PROTACs have been successful at degrading intracellular soluble proteins, their utility for degrading membrane proteins has not been fully explored.^15–16^ This is, in part, because membrane proteins are primarily degraded through lysosomal pathways such as receptor-mediated endocytosis and autophagy and may not be optimally accessible for PROTACs.^17-18^ In recent years, lysosome targeting chimeras (LYTACs), antibody-based PROTACs (AbTACs), and proteolysis targeting antibodies (PROTABs) have emerged as promising proximity-inducing modalities to target secreted and membrane proteins for lysosomal degradation, however to date such approaches have mainly relied on the use of antibodies.^19–24^ Therefore, we anticipate small molecule-based strategies that function intracellularly for the degradation of membrane proteins through lysosomal machineries could expand the scope of TPD modalities.

Lysosomal membrane proteins (LMPs) regulate essential cellular processes such as membrane repair, autophagy, phagocytosis, and viral infection.^25–26^ A recent study has demonstrated that a subset of LMPs, such as RNF152 and LAPTM4a, are short-lived and are internalized into the lysosome for degradation in a ubiquitin- and endosomal sorting complexes required for transport (ESCRT)-dependent manner, referred to as lysosomal microautophagy.^27–29^ We hypothesized that the rapid turnover and lysosomal proximity of LMPs could be co-opted to regulate the fate of proteins of interest (POIs) using proximity-inducing small molecules. We turned this concept into practice and have developed <u>Ly</u>sosome <u>M</u>embrane <u>Ta</u>rgeting <u>C</u>himeras (LYMTACs), heterobifunctional molecules composed of a POI ligand, a linker, and an LMP ligand (Figure 1A). Mechanistically, a LYMTAC induces the proximity between a POI and a short-lived LMP, thereby inducing relocalization of the target protein to the lysosomal membrane, promoting its internalization and subsequent degradation.

**Figure 1.**
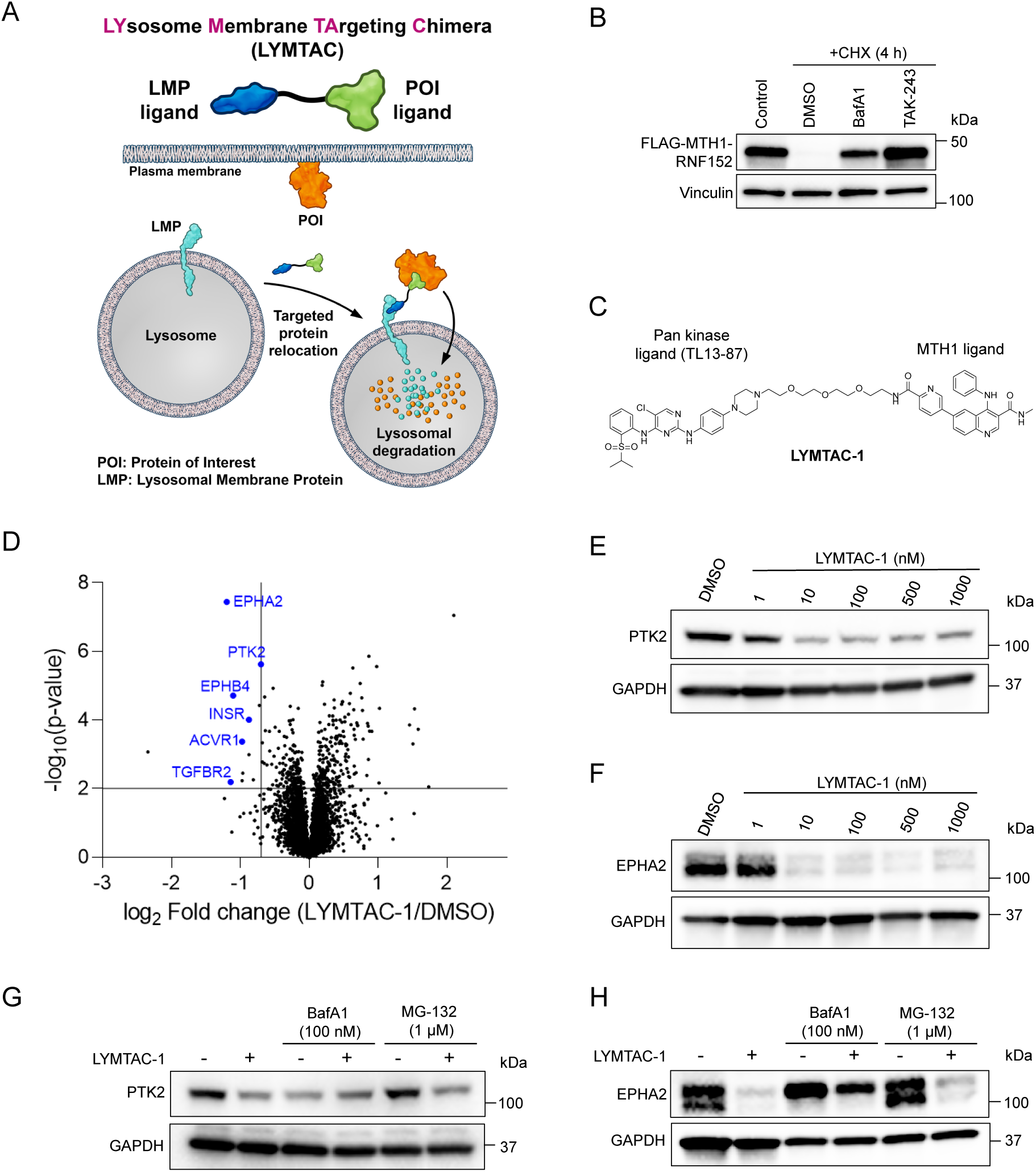
Development of LYMTAC as a modular tool for lysosomal degradation of membrane proteins. A) Schematic of the LYMTAC concept. A LYMTAC is a heterobifunctional molecule composed of a POI ligand, a linker and an LMP ligand. LYMTACs co-opt short-lived lysosomal membrane proteins as effectors to deliver target proteins for lysosomal degradation. B) Cycloheximide (CHX) assay in FLAG-MTH1-RNF152 stably expressing HCT116 cells. Cells were pre-treated with DMSO, 200 nM Bafilomycin A1 (BafA1), or 1 µM ubiquitin E1 inhibitor (TAK-243) for 30 min, followed by co-treatment with 100 µg/mL CHX for 4 h. Cells were harvested at 0 h to include as a control. Cells were lysed and subjected to immunoblotting with MTH1 and vinculin antibodies. Data are representative of two independent experiments. C) Structure of LYMTAC-1 (promiscuous kinase inhibitor-PEG2-MTH1 ligand). D) Quantitative proteomics analysis of LYMTAC-1. HEK293T cells stably expressing FLAG-MTH1-RNF152 were treated with either DMSO or 500 nM LYMTAC-1 for 19 h and subjected to global proteomics analysis. Data are representative of three treatment replicates. E) LYMTAC-1 induced dose-dependent degradation of PTK2. HEK293T cells stably expressing FLAG-MTH1-RNF152 were treated with increasing concentrations of LYMTAC-1 for 19 h and subjected to immunoblotting. Data are representative of three independent experiments. F) LYMTAC-1 induced dose dependent degradation of EPHA2. Data are representative of three independent experiments. G-H) HEK293T cells stably expressing MTH1-RNF152 were pre-treated with BafA1 and MG-132 for 30 min, co-treated with DMSO or 500 nM LYMTAC-1 for 19 h, and subjected to immunoblotting with the indicated antibodies. Data are representative of two independent experiments.

By applying this concept to various membrane kinases and oncogenic KRAS^G12D^, we demonstrate the utility of LYMTACs in targeting membrane proteins for lysosomal clearance. Furthermore, we demonstrate that LYMTACs are capable of exerting pharmacological activity through substrate relocalization and sequestration at the lysosome even in the absence of extensive degradation. This dual mode of action translated into a deeper suppression of KRAS^G12D^ signaling and inhibition of cancer cell viability compared to reversible KRAS inhibitors. The scope of this platform is further highlighted by establishing that multiple LMPs, such as RNF152, LAMPT4a, and LAPTM5, can serve as effectors for the LYMTAC technology. We anticipate that the chemical modularity and tunability of small molecule LYMTACs will offer new opportunities to optimize such agents for the efficient degradation of diverse membrane-associated proteins.

## Results and Discussion

### Development of LYMTAC as a modular tool for lysosomal degradation of membrane proteins

To establish the LYMTAC technology, we used RNF152 as our primary LMP effector for proof-of-concept studies. Due to the lack of selective and well-characterized ligands for RNF152, we generated a model system using MTH1-tagged RNF152 (MTH1-RNF152) to allow for induced proximity to the POI using a selective MTH1 binding ligand.^30^ Importantly, MTH1 tagging did not alter the turnover rate of RNF152 as shown by almost complete degradation of MTH1-RNF152 after only four hours of protein synthesis inhibition by cycloheximide in HEK293T and HCT116 cells (Figure 1B, Figure S1). Furthermore, this rapid turnover of MTH1-RNF152 was both ubiquitin- and lysosome-dependent as evidenced by inhibition by the ubiquitin E1 inhibitor (TAK-243) and lysosomal V-ATPase inhibitor (Bafilomycin A1) (Figure 1B).

After establishing the MTH1-RNF152 LMP model cellular system, we probed the impact of LYMTAC on target protein degradation. As such, we devised an untargeted approach using a promiscuous kinase inhibitor as the POI ligand.^31^ We generated a multi-kinase targeting LYMTAC (LYMTAC-1) by linking the promiscuous kinase inhibitor, TL-13-87, to an MTH1-targeting ligand (Figure 1C). This non-selective kinase inhibitor binds to a vast array of membrane, cytoplasmic, and nuclear protein kinases, thus allowing us to identify the target space that is most readily addressed by LYMTACs. HEK293T cells stably expressing MTH1-RNF152 were treated with LYMTAC-1 and subjected to global proteomics to identify degradable protein targets in an unbiased manner. A total of six protein kinases were significantly downregulated by LYMTAC-1, validating the ability of LYMTACs to drive target protein degradation (Figure 1D). Strikingly, the kinases that were downregulated by LYMTAC-1 were exclusively membrane proteins, including both integral (EPHA2, INSR, EPHB4,

ACVR1, TGFBR2) and peripheral (PTK2) membrane proteins. ^32–33^ Intriguingly, previously reported CRBN-and VHL-based pan-kinase PROTACs induced degradation of mostly cytosolic and nuclear kinases.^31,34^ While PTK2 was a common membrane-associated protein degraded by both modalities, the integral membrane proteins (EPHA2, INSR, EPHB4, ACVR1, TGFBR2) were not degraded by PROTACs. This is in stark contrast to LYMTAC-1 that mainly induced lysosomal degradation of membrane-associated protein kinases. Thus, LYMTACs provide access to a distinct pool of cellular targets that has been challenging to target using PROTACs.

To corroborate the proteomics results, HEK293T cells stably expressing MTH1-RNF152 were treated with increasing concentrations of LYMTAC-1 and subjected to immunoblotting. Consistent with the proteomics data, LYMTAC-1 induced PTK2 and EPHA2 degradation in a dose-dependent manner, with robust degradation observed at 10 nM (Figure 1E,1F). LYMTAC-1 activity was dependent on RNF152, since the addition of excess MTH1 ligand abrogated PTK2 degradation (Figure S2). Next, we investigated the route of degradation. Cells were pre-treated with either lysosome (Bafilomycin A1) or proteasome (MG-132) inhibitors, and LYMTAC-1-induced degradation of PTK2 and EPHA2 was assessed. LYMTAC-1-induced PTK2 and EPHA2 degradation was rescued by Bafilomycin A1 but not MG-132, consistent with lysosomal-mediated clearance of these membrane proteins, further confirming the anticipated degradation mechanism of LYMTACs (Figure 1G, 1H).

### LYMTACs induce ubiquitylation and lysosomal degradation of KRAS^G12D^

Having established the ability of LYMTACs to target membrane proteins for degradation, we sought to further investigate the therapeutic potential of these agents in degrading a historically challenging-to-drug protein, KRAS^G12D^. KRAS^G12D^ is a membrane-anchored protein that has been associated with tumorigenic capacity of many cancer types.^35^ We first utilized a chemical genetic system by co-expressing MTH1-RNF152 effector and HiBiT-FKBP12^F36V^-KRAS^G12D^ POI (Figure 2A).^36–37^ We chose to first establish this system using dimerization tags in cells not dependent on KRAS signaling for survival. Endogenously tagged HCT116 cells were generated by knocking in HiBiT-FKBP12^F36V^at the KRAS locus where gene editing was used to introduce G12D by correcting the G13D mutation that was present naturally in HCT116 cells. This approach allowed us to quantitatively assess KRAS^G12D^ levels and ligand engagement without the complications associated with the use of pharmacologically active inhibitors in cells sensitive to KRAS^G12D^ depletion. Next, we designed LYMTAC-2, comprising an FKBP12^F36V^ ligand, a connecting linker, and an MTH1 ligand to dimerize the target and effector proteins (Figure 2B). In line with our pan-kinase data, RNF152-recruiting LYMTAC-2 induced potent but partial KRAS^G12D^ degradation in a dose-dependent manner after 24 hours (Figure 2C).

**Figure 2.**
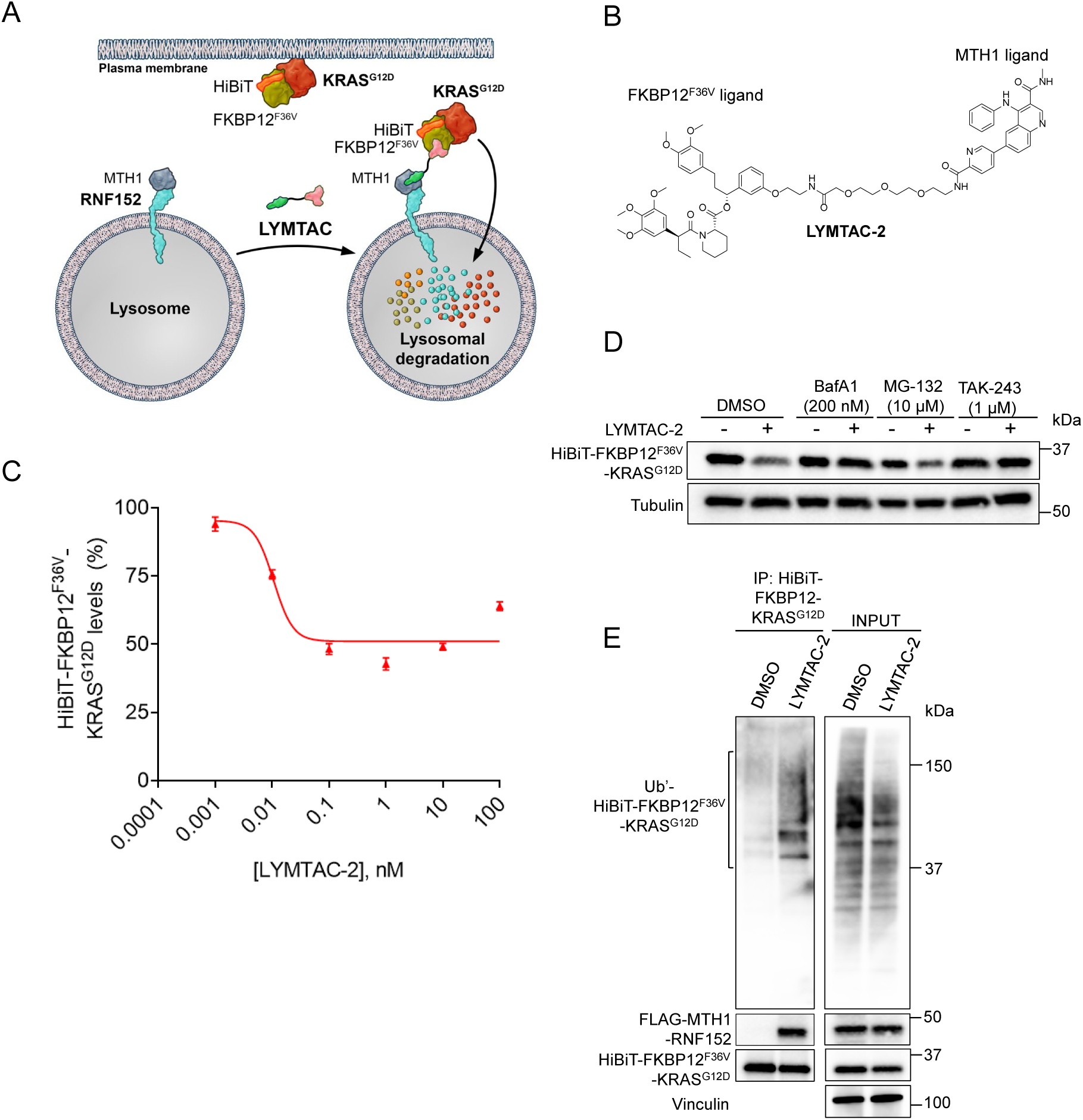
LYMTACs induce ubiquitylation and lysosomal degradation of KRAS^G12D^. A) Schematic of the chemical genetic system where the target protein is HiBiT-FKBP12^F36V^-KRAS^G12D^, and the effector is MTH1-RNF152. B) Structure of LYMTAC-2 (FKBP12^F36V^ ligand-PEG2-MTH1 ligand). C) HiBiT cellular degradation assay in MTH1-RNF152 stably expressing, knock-in HCT116 (HiBiT-FKBP12^F36V^-KRAS^G12D:^ (HF-KRAS)) cells. Cells were treated with increasing concentrations of LYMTAC-2 for 24 h and HiBiT levels were measured using Nano-Glo® HiBiT lytic reagent. Data are representative of three independent experiments, reported as the mean ± S.E. D) HCT116 (HF-KRAS) cells stably expressing FLAG-MTH1-RNF152 were pre-treated with BafA1, MG-132, or TAK-243 for 30 min followed by co-treatment with DMSO or 500 nM LYMTAC-2 for 6 h and subjected to immunoblotting with KRAS and tubulin antibodies. Data are representative of three independent experiments. E) KRAS^G12D^ ubiquitylation assay. HA-ubiquitin was transfected into HCT116, pre-treated with 200 nM BafA1 for 30 min, then co-treated with DMSO or 1 µM LYMTAC-2 for 4 h. Next, cells were lysed and subjected to immunoprecipitation with anti-HiBiT antibody. Whole cell lysate was used as INPUT. The respective blots are probed with HA, MTH1, HiBiT, and vinculin antibodies. Data are representative of three independent experiments.

To obtain additional insights on LYMTAC-2-induced KRAS^G12D^ degradation, HCT116 cells were pre-treated with inhibitors such as Bafilomycin A1, MG-132, or TAK-243, and then co-treated with LYMTAC-2 and subjected to immunoblotting. LYMTAC-2-mediated KRAS^G12D^ degradation was rescued by both ubiquitin E1 (TAK-243) and lysosomal (Bafilomycin A1) inhibitors (Figure 2D). However, LYMTAC-induced KRAS^G12D^ degradation was not rescued by proteasome inhibition. These data indicate that LYMTAC-2-mediated KRAS^G12D^ degradation follows the native ubiquitin- and lysosome-dependent turnover of RNF152. RNF152 is an E3 ubiquitin ligase LMP whose rapid internalization and lysosomal degradation is dependent on its own auto-ubiquitylation.^27^ Thus, we speculated that LYMTAC-2 may drive degradation of KRAS^G12D^ through RNF152-dependent ubiquitylation of KRAS^G12D^. To this end, we performed a cellular ubiquitylation assay by transiently expressing HA-ubiquitin in HCT116 cells followed by immunoprecipitation and immunoblotting. Notably, cells treated with LYMTAC-2 revealed rapid KRAS^G12D^ ubiquitylation (Figure 2E). Collectively, our data suggest that LYMTACs can lead to the formation of a productive ternary complex with KRAS^G12D^ and RNF152, promote KRAS^G12D^ ubiquitylation, and induce lysosome-dependent clearance of KRAS^G12D^.

### KRAS^G12D^ LYMTACs induce pathway suppression irrespective of KRAS^G12D^ degradation

Emerging resistance to KRAS inhibitors (KRASi) is a critical challenge in maximizing the clinical potential of these drugs.^38–41^ Degradation strategies that remove rather than inhibit oncogenic proteins offer the potential of addressing a number of such resistance mechanisms, which has encouraged our development of novel proximity-inducing approaches to degrade membrane-associated oncogenic KRAS. Having established and characterized the biochemical activity of KRAS^G12D^ LYMTACs, we next investigated the downstream consequences of lysosomal-mediated degradation of KRAS^G12D^. To target endogenous KRAS^G12D^ directly, we generated KRAS^G12D^ LYMTACs by replacing the FKBP12^F36V^ ligand with a pan-KRAS inhibitor (pan-KRASi) (Figure 3A, 3B). Extensive research in the TPD field has defined several common structural features that influence the efficacy of PROTAC design such as exit vector orientation, linker length, linker composition, and E3 ligase ligand.^42–44^ Borrowing from this knowledge, we sought to investigate how such parameters impacted LYMTAC efficacy. X-ray crystallographic characterization of the binding mode of the pan-KRAS inhibitor has suggested attachment of the effector ligand via the fused bicyclo-pyrollidine ring could yield active PROTACs and LYMTACs.

**Figure 3.**
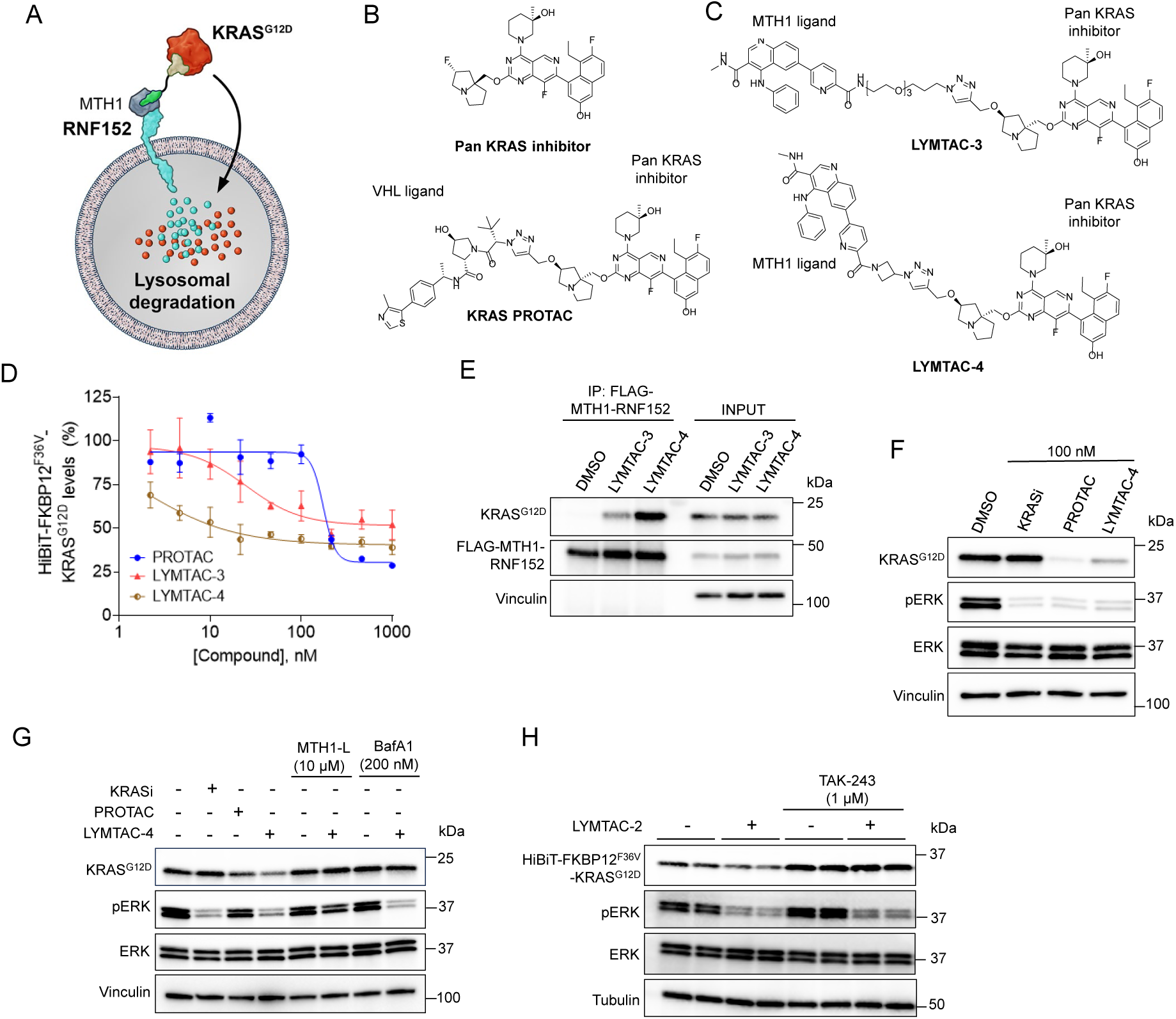
KRAS LYMTACs induce pathway suppression irrespective of KRAS degradation. A) Schematic of complex formation between KRAS^G12D^ and MTH1-RNF152 in the presence of KRAS-targeting LYMTAC. B) Structures of the pan-KRAS inhibitor and PROTAC C) Structures of LYMTAC-3 and LYMTAC-4. D) HiBiT cellular degradation assay in FLAG-MTH1-RNF152-stably expressing, knock-in HCT116 (HiBiT-FKBP12^F36V^-KRAS^G12D^) cells. Cells were treated with indicated doses of compounds for 24 h and subjected to Nano-Glo® HiBiT lytic assay. Data are representative of three independent experiments, reported as the mean ± S.E. E) Ternary complex formation assay, AsPC-1 cells stably expressing FLAG-MTH1-RNF152 were pre-treated with BafA1 for 30 min, followed by co-treatment with DMSO or 1 µM LYMTAC-3 or LYMTAC-4 for 4 h and subjected to immunoprecipitation with anti-FLAG antibody. Whole cell lysate was used as INPUT. The respective blots are probed with KRAS, MTH1, and vinculin antibodies. Data are representative of two independent experiments. F) AsPC-1 cells stably expressing FLAG-MTH1-RNF152 were treated with DMSO or 100 nM of the indicated compounds for 24 h and subjected to immunoblotting with KRAS, pERK, ERK, and vinculin antibodies. Data are representative of three independent experiments. G) AsPC-1 cells stably expressing FLAG-MTH1-RNF152 were treated with 100 nM KRAS inhibitor, 100 nM PROTAC, or 100 nM LYMTAC-4 for 6 h. For control experiments with LYMTAC-4, cells were pretreated with MTH1-ligand or BafA1 for 30 min, followed by co-treatment in the absence or presence of 100 nM LYMTAC-4 for 6 h, and subjected to immunoblotting with KRAS, p-ERK, ERK, and vinculin antibodies. Data are representative of two independent experiments. H) HEK293T cells stably expressing HiBiT-FKBP12-KRAS^G12D^ and FLAG-MTH1-RNF152 were pre-treated with 1 µM TAK-243 for 30 min followed by DMSO or 1 µM LYMTAC-2 treatment for 6 h. The respective blots are probed with KRAS, p-ERK, ERK, and tubulin antibodies. Data are representative of two independent experiments.

First, we screened LYMTACs with various PEG and rigid linkers for cellular HiBiT-FKBP12^F36V^-KRAS^G12D^ degradation using HCT116 cells and compared their efficacy to that achieved through PROTAC-mediated degradation (Figure 3B and 3C). While our KRAS PROTAC and LYMTACs displayed varying levels of KRAS^G12D^ degradation, rigid linker-based LYMTAC-4 (which also tethered the MTH1 ligand to the pan-KRAS inhibitor via a shorter linker) displayed improved D_max_ and DC_50_ values compared to PEG linker-based LYMTAC-3 (Figure 3D). Leveraging the more potent LYMTAC-4 in subsequent experiments, we functionally characterized the effect of LYMTAC-induced KRAS^G12D^ degradation in AsPC-1 cells, a relevant KRAS^G12D^ mutant pancreatic cancer cell line that relies on this oncogene for cell proliferation and survival.^45^ MTH1-RNF152 was stably introduced into AsPC-1 cells to evaluate ternary complex formation (TCF) by various LYMTACs. The data suggest that LYMTAC-4 induced a more productive ternary complex between KRAS^G12D^ and RNF152, compared to LYMTAC-3 (Figure 3E). Experience with this system further supported the selection of LYMTAC-4 as an appropriate tool molecule for subsequent investigation. To evaluate the cellular effect of LYMTAC-4, AsPC-1 cells were treated with pan-KRASi, PROTAC, or LYMTAC-4 for 24 hours and probed for pathway activity. LYMTAC-4 induced KRAS degradation and robust pERK downregulation at 24 hours, comparable to pan-KRASi and PROTAC (Figure 3F).

We next sought to further elucidate LYMTAC mode of action and its dependence on TCF. This was investigated by treating AsPC-1 cells with LYMTAC-4 for 6 hours in the absence or presence of excessive competing MTH1-ligand or Bafilomycin A1. Excess MTH1 ligand abrogated LYMTAC-induced KRAS^G12D^ degradation and p-ERK signaling, demonstrating that TCF is required to induce KRAS^G12D^ degradation and inhibit downstream signaling (Figure 3G). As expected, Bafilomycin A1 treatment rescued LYMTAC-4-induced KRAS^G12D^ degradation. Unexpectedly, in Bafilomycin A1-treated cells, LYMTAC-4 still suppressed KRAS^G12D^ downstream signaling even in the absence of KRAS^G12D^ degradation. To rule out that p-ERK downregulation was due to the pan-KRASi moiety of LYMTAC-4, we stably co-expressed FKBP12^F36V^-KRAS^G12D^ and MTH1-RNF152 in HEK293T cells and engaged FKBP12-KRAS with a non-inhibitory FKBP12 ligand containing LYMTAC-2 (FKBP12 ligand-PEG2-MTH1 ligand). Strikingly, FKBP12-targeting LYMTAC-2 also suppressed p-ERK signaling irrespective of KRAS ubiquitylation and degradation (Figure 3H). Collectively, these data provide compelling evidence that pathway inhibition is not due to the inhibitory effect of pan-KRASi-based LYMTACs but could be attributed to the ability of LYMTACs to affect target biology beyond target degradation. Overall, the data indicate that LYMTAC-induced KRAS ubiquitylation and degradation are not uniformly required for downstream signaling inhibition, while TCF is critical to suppress p-ERK signaling.

### KRAS relocalization and lysosomal degradation using LYMTAC lead to deeper pathway suppression and potent cell killing compared to SMI

The observation that signaling downstream of KRAS^G12D^ can be modulated by LYMTACs even in the absence of KRAS^G12D^ degradation prompted us to further explore whether quantitative relocalization of KRAS from the plasma membrane to the lysosome could be driving this effect (Figure 4A). To address this hypothesis, we generated mNeonGreen-KRAS^WT^ stably expressing HEK293T cells to allow for subcellular KRAS tracking by confocal microscopy. Pre-treatment of these cells with Bafilomycin A1 followed by LYMTAC-4 revealed rapid and marked KRAS relocalization from the plasma membrane followed by evident association with RNF152 at the lysosome, as evidenced by co-localization with the lysosome marker LAMP1 (Figure 4B). This was specific for LYMTACs as this phenotype was absent in cells treated with pan-KRASi or PROTAC (Figure 4C).

**Figure 4.**
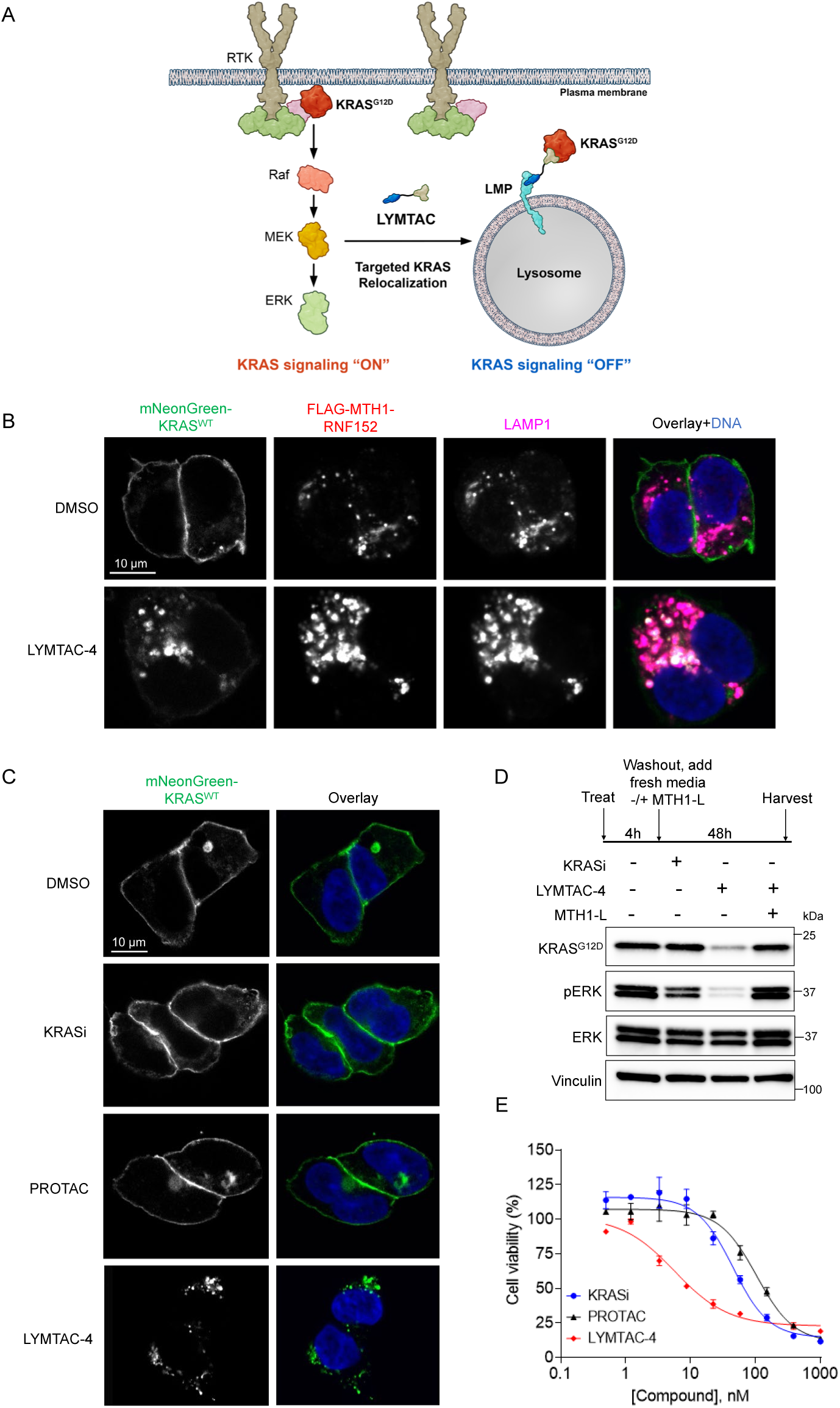
KRAS relocalization and lysosomal degradation using LYMTAC lead to deeper pathway suppression and potent cell killing compared to SMI. A) Schematic of LYMTAC-induced KRAS relocalization from plasma membrane to lysosome. B) KRAS localization analyzed by confocal microscopy. FLAG-MTH1-RNF152-stably expressing HEK293T mNeonGreen-KRAS ^WT^ cells were pre-treated with 200 nM BafA1 for 30 min, and then co-treated with DMSO or 1 µM LYMTAC-4 for 4 h. Cells were fixed, permeabilized, and immunostained with FLAG and LAMP1 antibodies, followed by appropriate secondary fluorophore-conjugated antibodies. Nuclei were stained with Hoechst dye. Data are representative of two independent experiments. C) Confocal images in FLAG-MTH1-RNF152 stably expressing HEK293T mNeonGreen-KRAS ^WT^ cells were pre-treated with 200 nM BafA1 or 1 µM Bortezomib for 30 min before 4 h co-treatment with DMSO (+BafA1), 1 µM pan-KRAS inhibitor, 1 µM PROTAC (+Bortezomib), or 1 µM LYMTAC-4 (+BafA1). Cells were fixed and nuclei were stained with Hoechst dye. Data are representative of two independent experiments. D) Washout experiment in AsPC-1 cells stably expressing FLAG-MTH1-RNF152. Cells were treated with 100 nM pan-KRASi and 100 nM LYMTAC-4 for 4 h, washed three times with PBS and replaced with fresh media in the absence or presence of 5 µM MTH1-ligand for 48 h. Data are representative of two independent experiments. E) pan-KRASi, PROTAC, and LYMTAC-4 activity in a 5-day cell proliferation assay in AsPC-1 cells stably expressing MTH1-RNF152. Data are representative of four independent experiments, reported as the mean ± S.E.

KRAS is a plasma membrane anchored protein that activates oncogenic signaling at the membrane via multiple effector proteins.^46^ Various attempts to block KRAS trafficking to the plasma membrane have been made, yet these approaches have been largely unsuccessful.^47–48^ In addition, membrane-bound KRAS is susceptible to pathway rebound driven by upstream receptor tyrosine kinases such as EGFR, ultimately contributing to therapeutic resistance.^40,49–50^ Given that LYMTACs display dual functionality in the inactivation of membrane proteins (i.e., via intracellular relocalization and via subsequent lysosomal degradation), their distinct mechanism of action may offer multiple advantages in overcoming some of the limitations commonly seen with current therapies. We explored this possibility by measuring pathway rebound upon drug withdrawal. AsPC-1 cells were treated with pan-KRASi or LYMTAC-4 for four hours, excess compound was washed away, and fresh media was added for 48 hours before analysis. Consistent with the reported KRASi-induced signaling rebound, we observed a p-ERK signal rebound after 48 hours in the pan-KRASi-treated samples (Figure 4D). Strikingly, LYMTAC-4 induced sustained KRAS^G12D^ degradation and delayed rebound of p-ERK signaling compared to pan-KRASi (Figure 4D). Excess MTH1 ligand rescued KRAS^G12D^ degradation and p-ERK inhibition, indicating that this effect was on target and dependent on RNF152 (Figure 4D). Given that LYMTAC-4 maintained prolonged signaling inhibition, we sought to further monitor the effect of LYMTACs on AsPC-1 cell proliferation. Importantly, the depth of pathway suppression also translated phenotypically as LYMTAC-4 demonstrated improved cell killing compared to both KRASi and PROTAC (Figure 4E). This potent cell killing could be attributed to the multi-pharmacology displayed by LYMTACs via KRAS relocalization, sustained p-ERK inhibition, and lysosomal degradation, which sets LYMTACs apart from existing modalities.

### Expansion of the LYMTAC platform to other LMPs

To demonstrate the generality of our platform, we targeted additional LMP effectors beyond RNF152. We focused on LAMPT4a and LAPTM5, two LMPs that associate with the NEDD4-1 E3 ubiquitin ligase at the lysosome membrane through PY motifs (Figure 5A).^27, 29^ After generating cells stably expressing MTH1-tagged LAPTM4a and LAPTM5, we first performed a cycloheximide assay in HEK293T cells to confirm their short-lived nature. The data indicate that both MTH1-LAPTM4a and MTH1-LAPTM5 are rapidly degraded within four hours in HEK293T cells (Figure S3). Similarly to HEK293T cells, HCT116 cells stably expressing MTH1-tagged LAPTM4a or LAPTM5 showed rapid LMP turnover in a ubiquitin and lysosome-dependent manner (Figure 5B and 5C).

**Figure 5.**
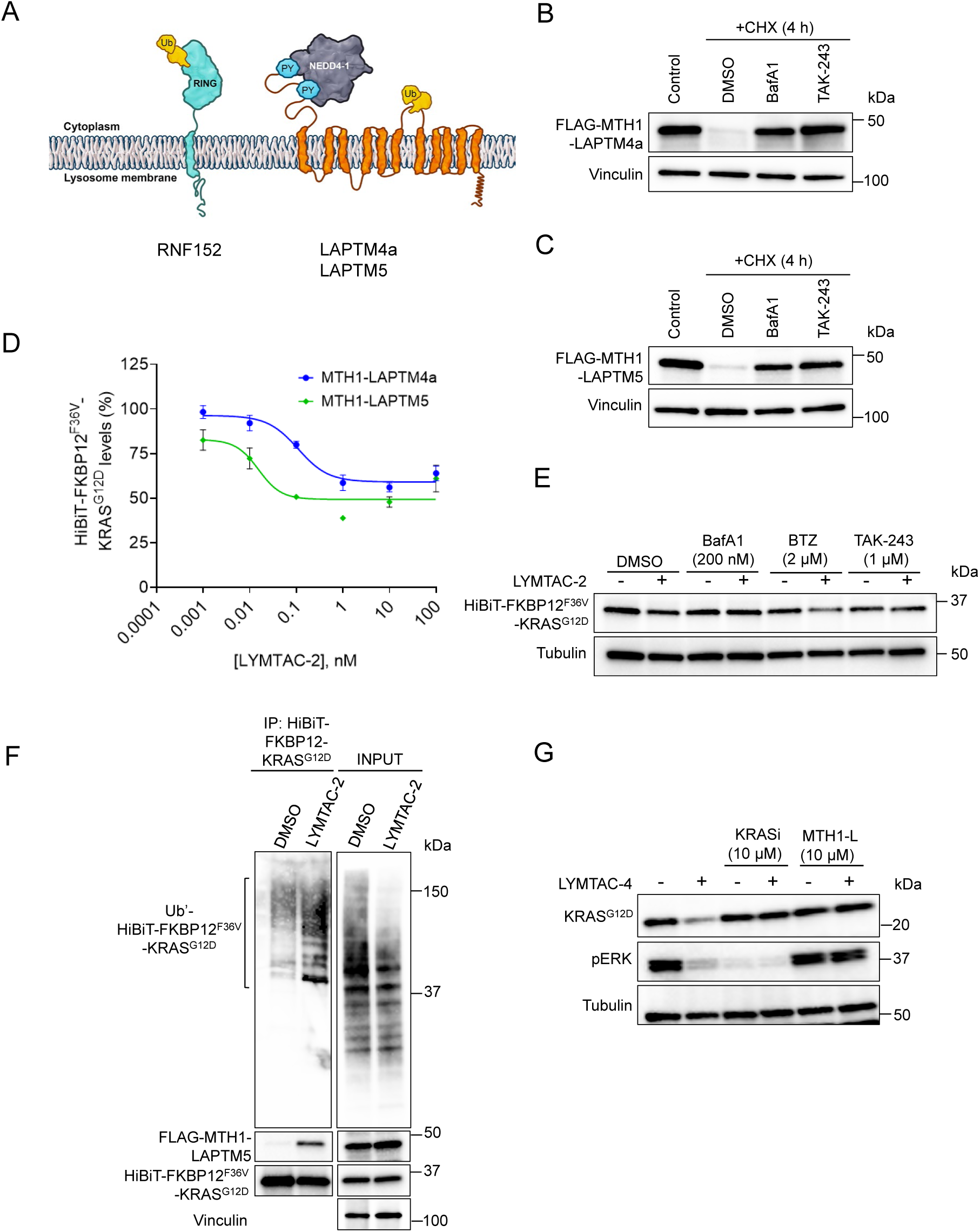
Expansion of LYMTAC platform to other LMPs. A) Schematic of Lysosome Membrane Proteins (LMPs). B) CHX assay in FLAG-MTH1-LAPTM4a-stably expressing HCT116 cells. Cells were pre-treated with DMSO, 200 nM Bafilomycin A1 (BafA1), or 1 µM E1 inhibitor (TAK-243) for 30 min, followed by co-treatment with CHX for 4 h. Cells were harvested at 0 h to include as a control. Cells were lysed and subjected to immunoblotting with MTH1 and vinculin antibodies. Data are representative of three independent experiments. C) CHX assay in FLAG-MTH1-LAPTM5-stably expressing HCT116 cells. Data are representative of three independent experiments. D) HiBiT cellular degradation assay in FLAG-MTH1-LAPTM4a- and FLAG-MTH1-LAPTM5-stably expressing HCT116 (knock-in-HiBiT-FKBP12^F36V^-KRAS^G12D:^ (HF-KRAS)) cells. Cells were treated with increasing concentrations of LYMTAC-2 for 24 h and HiBiT levels were measured using Nano-Glo® HiBiT lytic reagent. Data are representative of two independent experiments and reported as the mean ± S.E. E) HCT116 (HF-KRAS) cells stably expressing FLAG-MTH1-LAPTM5 were pre-treated with BafA1, 2 BTZ, or TAK-243 for 30 min followed by co-treatment with DMSO or 500 nM LYMTAC-2 for 6 h and subjected to immunoblotting with KRAS and tubulin antibodies. Data are representative of two independent experiments. F) KRAS ubiquitylation assay. HA-ubiquitin was transfected into HCT116 expressing FLAG-MTH1-LAPTM5, which were pre-treated with 200 nM BafA1 for 30 min followed by co-treatment with DMSO or 1 µM LYMTAC-2 treatment for 4 h. Next, cells were lysed and subjected to immunoprecipitation with anti-HiBiT antibody. Whole cell lysate was used as INPUT. The respective blots were probed with HA, MTH1, HiBiT, and vinculin antibodies. Data are representative of two independent experiments. G) AsPC-1 cells stably expressing FLAG-MTH1-LAPTM4a were pre-treated with either pan-KRAS inhibitor or MTH1 ligand, followed by DMSO or 100 nM LYMTAC-4 treatment for 6 h. Cells were lysed and subjected to immunoblotting with KRAS, pERK, and tubulin antibodies. Data are representative of two independent experiments.

Having confirmed the fast turnover of MTH1-LAPTM4a and MTH1-LAPTM5, we next asked whether these two LMPs can induce target protein degradation in a LYMTAC-dependent manner. To this end, we utilized HiBit-FKBP12^F36V^-KRAS^G12D^ knock in HCT116 cells and stably expressed MTH1-LAPTM4a or MTH1-LAPTM5. Consistent with results seen for MTH1-RNF152, both MTH1-LAPTM4a and MTH1-LAPTM5 induced potent KRAS^G12D^ degradation in a LYMTAC-2-dependent manner (Figure 5D), validating these two LMPs as effectors for the LYMTAC technology. To verify the mechanism of action for LAPTM5-based LYMTACs, HCT116 cells were treated with lysosomal (Bafilomycin A1), proteasomal (bortezomib), and ubiquitin E1 (TAK-243) inhibitors prior to LYMTAC-2 treatment and subjected to immunoblotting. Consistent with RNF152-recruiting LYMTACs, LAPTM5-induced KRAS^G12D^ degradation was rescued by both lysosome and ubiquitin E1 inhibitors, but not proteosome inhibitor (Figure 5E). To further understand the ubiquitin-dependent mechanism of action, cellular ubiquitylation assays were performed. As expected, LAPTM5 formed a ternary complex with KRAS^G12D^ and induced KRAS^G12D^ ubiquitylation in the presence of LYMTAC-2 (Figure 5F).

Finally, to expand LMPs that can be harnessed for LYMTAC development, we generated MTH1-LAPTM4a expressing AsPC-1 cells and confirmed the rapid degradation of this construct as had been observed with prior LMP (Figure S4). We further confirmed that KRAS^G12D^ levels were unaffected by overexpressing either MTH1-RNF152 or MTH1-LAPTM4a in the absence of LYMTAC molecules (Figure S4). Similarly to RNF152, LAPTM4a formed a productive ternary complex with KRAS^G12D^ in the presence of LYMTAC-4 (Figure S5). Moreover, LAPTM4a-recruiting LYMTAC-4 induced KRAS^G12D^ degradation and p-ERK inhibition in AsPC-1 cells without affecting the levels of MTH1-LAPTM4a (Figure 5G, Figure S6). To further verify the mechanism of action, cells were treated with either pan-KRASi or MTH1-ligand prior to LYMTAC-4 treatment. The ligand competition data indicate that LYMTAC-induced KRAS^G12D^ degradation and p-ERK inhibition are LAPTM4a-dependent and require a ternary complex between KRAS^G12D^ and LAPTM4a.

In summary, we describe here a SM-based LYMTAC platform that functions by repurposing short-lived lysosomal membrane protein effectors to induce membrane protein relocalization and subsequent degradation. Using a promiscuous kinase inhibitor-based LYMTAC, we demonstrate that LYMTACs preferentially deliver membrane kinases for lysosomal degradation. The observed substrate selectivity was a differentiating feature of LYMTAC-driven degradation when compared to cytosolic and nuclear protein degradation by a promiscuous kinase inhibitor-based PROTAC. The selectivity of LYMTACs towards membrane-associated proteins is likely explained by their natural turn over in the lysosome and continuous flux in the membrane trafficking pathways that offer opportunities for these membrane proteins to be captured by LMPs. Furthermore, using oncogenic KRAS^G12D^ as a therapeutically relevant, membrane-associated target, we demonstrated that LYMTACs induce sustained pathway inhibition and potent cell killing as compared to a KRAS inhibitor. Notably, LYMTAC-induced KRAS relocalization from the plasma membrane to the lysosome is sufficient to suppress downstream signaling. While we observe lack of complete KRAS degradation (D_max_ ∼50-60%) with LYMTACs, we observe quantitative relocalization of KRAS from plasma membrane to lysosomes, potent inhibition of p-ERK signaling, and cytotoxicity.

Our proof-of-concept results demonstrate that the LYMTAC technology could be an effective strategy in dampening KRAS inhibitor resistance mechanisms by sequestrating KRAS away from the plasma membrane. This is reminiscent of proximity-inducing modalities that have recently been shown to modulate target protein via non-degradative mechanisms such as relocalization and sequestration.^51–55^ In addition to the in-depth characterization of RNF152, we validated the LYMTAC technology with other LMPs such as LAPTM4a and LAPTM5, clearly demonstrating the potential compatibility of many more LMPs with LYMTAC technology. Overall, LYMTACs are a small molecule proximity-based platform that function intracellularly to inhibit oncogenic signaling through multi-pharmacology, including relocalization (occupancy-driven pharmacology) and lysosomal degradation (event-driven pharmacology) of membrane proteins (Figure 6). While we introduce LYMTAC as a chemical biology-driven proof-of-concept study, these MTH1-derived LYMTAC tool molecules can be utilized in fundamental research to functionally characterize the effect of target protein relocalization and lysosomal degradation. Future work will focus on understanding the target scope, LMP space (including cell-type specificity), and ligand discovery for LMPs to enable the LYMTAC technology for therapeutic development.

**Figure 6.**
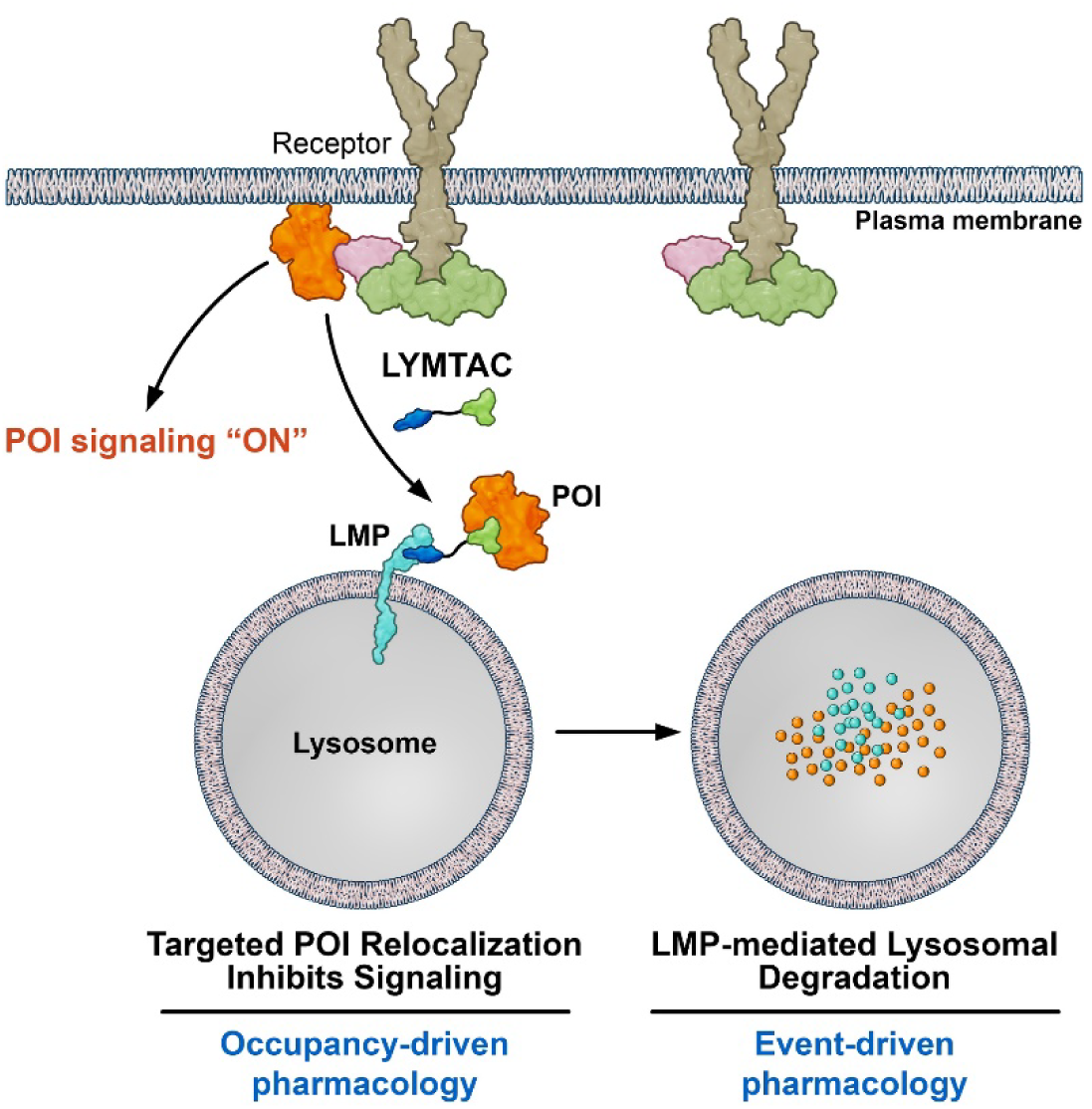
LYMTACs display multi-pharmacology. LYMTAC is a SM-based, proximity-inducing modality that functions intracellularly to relocalize and degrade membrane proteins via the lysosome. LYMTACs inhibit POI signaling via relocalization and subsequent lysosomal degradation.

## Acknowledgements

We thank Ryan Case, Jan Andersson, Olivier Bedel, Rati Verma, Kusal Samarasinghe, Spencer Hill, and Kyle Mangano for helpful discussions. We thank Craig Kiefer for generating graphics for the manuscript. We thank all members of IPP for their support.

## Author Contributions

D.A.N. and P.R.P. conceived the project and designed experiments. D. A.N., W.D., J.X., S.L., R.G.G., C.E.S., performed experiments and analyzed data. K.S.A., G.H., G.M., A.T.T., R.P.W., B.A.L., S.A., M.K.A., A.S., and R.K., designed and synthesized compounds. D.A.N. wrote the original draft of the manuscript and was revised with input from all authors.

## Materials and Methods

### Cell lines

Human embryonic kidney 293T (HEK293T), AsPC1, and HCT116 cells were obtained from ATCC. HEK293T cells were cultured in DMEM (Gibco) supplemented with 10% (v/v) heat-inactivated fetal bovine serum (FBS) and 1% penicillin/streptomycin (Pen/strep) (v/v). AsPC-1 and HCT116 cells were grown in RPMI media supplemented with 10% FBS and 1% Pen/strep. Jump-In™GripTite™ HEK293 cells (A14150) were obtained from Invitrogen and grown in DMEM Glutamax supplemented with 10% FBS, 0.1 mM NEAA, 25 mM HEPES, 1x Pen/strep, 600 µg/mL Geneticin. Cells were cultured at 37 °C and 5% CO_2_ in a humidified incubator and regularly screened for mycoplasma contamination.

### Antibodies and reagents

Antibodies against MTH1 (43918), FLAG (14793), Vinculin (13901), Tubulin (2144), pERK (9101), ERK (4695), PTK2 (3285), EPHA2 (6997), HA (3724), LAMP1(15665), and GAPDH (2118) were purchased from Cell Signaling Technology. HiBiT antibody (N7200) was obtained from Promega and KRAS antibody (LS-C175665) was purchased from LSBio. Alexa Fluor 568 (A11011) and Alexa Fluor 647 (A32728) conjugated secondary antibodies and Hoechst 33342 stain (H3570) were purchased from Invitrogen. Amersham ECL Rabbit IgG, HRP-linked whole secondary Ab (from donkey) (NA934) and Amersham ECL Mouse IgG, HRP-linked whole secondary Ab (from sheep) (NA931) were purchased from Cytiva. RIPA Lysis and Extraction Buffer (89900) and IP-lysis buffer (87787) was purchased from ThermoScientific. Protease inhibitor cocktail (11873580001) was obtained from Roche and phosphatase inhibitor cocktail (A32957) were obtained from ThermoScientific. 1,10-phenanthroline monohydrate (S5543), N-ethylmaleimide (S3692), and M3814 (S8586) were purchased from Selleckchem. PR-619 (4482) was purchased from Tocris. Nano-Glo® HiBiT lytic detection reagent (N3040) was purchased from Promega and CyQUANT™ direct cell proliferation assay (C35011) was obtained from Invitrogen. Bafilomycin A1 (54645), MG-132 (2194) and Cycloheximide (2112) were purchased from Cell Signaling Technology. TAK-243 (HY-100487) and Bortezomib (HY-10227) were obtained from MedChemExpress. Puromycin (ant-pr-1), blasticidin (ant-bl-05), and Geneticin (ant-gn-1) were purchased from InvivoGen. Pierce™ Anti-DYKDDDDK (FLAG) Magnetic Agarose beads (A36797) and 16% Formaldehyde (28906) were purchased from ThermoScientific. Protein G Magnetic Sepharose beads (28951379) were obtained from Cytiva. 8 well Ibidi dishes (80826) were purchased from Ibidi and 4X NuPAGE™ LDS sample buffer (NP0007) was purchased from Invitrogen.

### Plasmids

HA-ubiquitin cDNA in pcDNA3.1+ vector was obtained from GenScript. FLAG-MTH1-RNF152, FLAG-MTH1-LAPTM4a, and FLAG-MTH1-LAPTM5 cDNA were cloned into pGenLenti-puro vector (GenScript). Lentivirus particles were used to transduce HEK293T, HCT116, and AsPC-1 cells and generated stable cells expressing different MTH1 fusions.

### Generation of endogenously tagged KRAS^G12D^(HiBiT-FKBP12^F36V^-KRAS^G12D^) in HCT116 cells

HCT116 cells have a heterozygous G13D mutation on the endogenous KRAS locus. To generate HCT116 cells with N-terminal HiBiT-FKBP12^F36V-^KRAS^G12D^ homozygous clones, cutting site was designed near the endogenous KRAS G12 locus (guide RNA: 5’-aaacttgtggtagttggagc-3’). The HDR template was designed to contain the BSD-P2A-HiBiT-FKBP12^F36V^ insertion and the G12D mutation simultaneously and de novo synthesized at GenScript. To generate the homozygous clones, HCT116 were seeded overnight in 12-well plates at 6×10^5^ cells per well overnight. The next day, cells were transfected with a mixture of 4x plasmids containing guide RNA, CRISPR enzyme, HDR template, and i53 HDR enhancer.^56^ Six hours after transfection, cells were passaged 1:8 into 6 well plates with 2.5 mL culture medium with 1 μM M3814 as another HDR enhancer. One day later, the media was replaced with media containing 2 μg/mL puromycin (the guide RNA plasmid contains a puromycin resistant gene for selection) and 1 μM M3814 for another 2 days. The media was then replaced with media containing 20 μg/mL blasticidin to select cells with endogenous KRAS tagging for 2 days. The cell pool was further recovered in regular medium for 2 days and were used for single cloning. For single cloning, approximately 500 cells were plated on a 10cm dish in tissue culture media and allowed to grow for approximately one week. Single colonies were isolated and transferred to a 96-well plate using a stereomicroscope and 20 μL pipette tips. Single clones were identified with genomic PCR and subjected to Sanger sequencing to confirm homozygous HiBiT-FKBP12^F36V^ tagging and G12D mutation at the N-terminus of endogenous KRAS.

### Generation of 293T, HCT116 (KI HiBiT-FKBP12-KRAS), and AsPC-1 cells stably expressing MTH1 fusions

To generate RNF152 expressing cells for global proteomics analysis, HEK293T cells were transduced with lentivirus particles containing FLAG-MTH1-RNF152 in the presence of polybrene (10 µg/mL). The cells were selected using 2 µg/ml puromycin for 2 weeks to generate stable cells and MTH1-RNF152 expression was verified using immunoblotting. FLAG-MTH1-LAPTM4a, and FLAG-MTH1-LAPTM5 stably expressing HEK293T cells were generated similar to RNF152. To generate effector expressing HCT116 (KI HiBiT-FKBP12^F36V-^ KRAS^G12D^) cells, cells were separately transduced with lentivirus particles containing FLAG-MTH1-RNF152, FLAG-MTH1-LAPTM4a, and FLAG-MTH1-LAPTM5, selected using 2 µg/mL puromycin for 2 weeks and verified stable expression via immunoblotting. To generate effector expressing AsPC-1 cells, cells were transduced with lentivirus particles containing FLAG-MTH1-RNF152 or FLAG-MTH1-LAPTM4a, selected using 1.5 µg/mL puromycin for 2 weeks and effector expression was verified via immunoblotting.

### Generation of HEK293 cells stably co-expressing HiBiT-FKBP12^F36V^-KRAS^G12D^ and MTH1-RNF152 fusion

To generate HiBiT-FKBP12^F36V-^KRAS^G12D^ stably expressing Jump-In™GripTite™ cells, HiBiT-FKBP12^F36V-^KRAS^G12D^ cDNA was cloned into pJTI™ R4 Dest CMV pA vector. This plasmid was co-transfected with pJTI™ R4 Int vector into Jump-In™GripTite™ HEK293 cells. After transfection, cells were selected using 10 µg/mL blasticidin to obtain single cell clones. HiBiT-FKBP12^F36V-^KRAS^G12D^ expression was verified by immunoblotting and HiBit assay. To generate effector expressing cells, HiBiT-FKBP12^F36V-^KRAS^G12D^ stable cells were transduced with lentivirus particles containing FLAG-MTH1-RNF152 in the presence of 10 µg/mL polybrene. The cells were selected using 2 µg/mL puromycin for 2 weeks to generate stable cells and MTH1-RNF152 expression was verified using immunoblotting.

### Generation of HEK293 cells stably co-expressing mNeonGreen-KRAS^WT^ and MTH1-RNF152

To generate mNeonGreen-KRAS^WT^ stable cells, mNeonGreen-KRAS^WT^ cDNA was cloned into pJTI™ R4 Dest CMV pA vector. This plasmid was co-transfected with pJTI™ R4 Int vector into Jump-In™GripTite™ HEK293 cells. After transfection, cells were selected for two weeks using 5 µg/mL blasticidin, and clones were obtained by single-cell sorting. mNeonGreen-KRAS^WT^ expression was verified by immunoblotting and microcopy analysis. To generate effector expressing cells, mNeonGreen-KRAS cells were transduced with lentivirus particles containing FLAG-MTH1-RNF152 in the presence of 10 µg/mL polybrene. The cells were selected using 2 µg/mL puromycin for 2 weeks to generate stable cells and MTH1-RNF152 expression was verified using immunoblotting.

### Cell lysis and Immunoblotting

After desired treatments, cells were washed 2X with PBS, trypsinized, and harvested as cell pellets. Cell pellets were lysed in RIPA buffer supplemented with 1X protease and 1X phosphatase inhibitor cocktail, incubated on ice for 20min and cell lysate were clarified by centrifugation. Lysate was then mixed with 4X NuPAGE™ LDS Sample Buffer supplemented with 10% β-mercaptoethanol, boiled at 95 °C, and proteins were separated on 4–20% Criterion TGX precast gradient gels (Bio-Rad) and transferred to PVDF membrane. Next, the blots were incubated with blocking buffer (5% (w/v) nonfat dry milk in TBST (0.1% Tween-20, 20 mM Tris–HCl pH 7.6, 150 mM NaCl) for 1 h at room temperature followed by overnight incubation with primary antibodies at 4 °C. After primary antibody incubation, membranes were washed 3X with TBST and incubated with HRP-conjugated secondary antibodies for 1 h at room temperature. Membrane was washed 3X with TBST, developed using ECL or Super Signal West Femto substrate (ThermoFisher), and images were acquired using Bio-Rad ChemiDoc system.

### Lysosome membrane protein (LMP) turnover analysis using cycloheximide assays

HCT116 cells stably expressing MTH1-LMP were seeded at a density of 6×10^5^ cells per well in a 6 well dish. After 24 h, cells were pre-treated with DMSO, 200 nM Bafilomycin A1, and 1 µM E1 inhibitor (TAK-243) for 30 min, followed by co-treatment with 100 µg/mL cycloheximide for 4 h. Cells were harvested at 0 h to include as a control. After 4h, cells were harvested and lysed with RIPA buffer supplemented with protease inhibitor cocktail and subjected to immunoblotting with indicated antibodies.

### Quantitative proteomics analysis

HEK293T cells stably expressing MTH1-RNF152 were seeded at a density of 5×10^5^ cells per well in a 6 well dish. After 24 h, cells were treated with DMSO and 500 nM LYMTAC-1 for 19 h. Next, cells were washed 1X with PBS, trypsinized, and harvested. Cell pellet was washed 2X with PBS and subjected to global proteomics analysis.

#### LC-MS sample preparation

Cell pellets were lysed in 100 µL of lysis buffer (2% SDS, 100 mM Tris-HCl) and sonicated using an Abcam PIXUL sonicator (50Hz, 20 min). The protein concentration of the lysates was determined by the BCA method (Thermo Fischer Scientific 23225). For each sample, 30 µg of protein was reduced and alkylated by adding Bond-Breaker TCEP (Thermo 77720) to a final concentration of 100 mM and chloroacetamide to a final concentration of 500 mM and incubating at 56 °C for 45 min. The samples were then subjected to the SP3 protocol for clean-up. Briefly, 30 µg of protein was bound to >300 µg of beads in 80% (v/v) ethanol for 15 min, and then washed three times with 80% ethanol and once with acetonitrile. The protein-bound beads were digested with 1.2 µg of Trypsin/Lys-C mix in 50 mM TEAB buffer for 18 h at 37 °C. The beads were removed, and the peptides were dried to eliminate TEAB. Before LC-MS analysis, the samples were resuspended in 5% formic acid.

#### LC-MS analysis method

Liquid Chromatography (LC) Conditions: The chromatographic separation was performed using a Thermo Scientific Vanquish Neo LC system with a Bruker PepSep C18 analytical column (15 cm x 150 µm, 1.5 µm particle size). The mobile phases were Solvent A (0.1% formic acid) and Solvent B (ACN80, acetonitrile with 20% water). The flow rate was set to 1.2 µL/min, and the column was equilibrated with 0% Solvent B before sample injection. The gradient elution was programmed as follows: 0% Solvent B from 0 to 1 minute, followed by an increase to 2% Solvent B at 1 minute. Subsequently, Solvent B was linearly ramped up to 24% over the course of 84 minutes (1-85 minutes). After reaching 24%, the percentage of Solvent B was further increased to 40% over the next 14 minutes (85-99 minutes). Finally, Solvent B was rapidly increased to 95% from 99 to 100 minutes to wash the column and held at 95% for 4 minutes (100-104 minutes) before the flow was returned to the initial conditions. Column re-equilibration was conducted at 95% Solvent B for an additional 4 minutes. The total run time for the method was 104 minutes, including the gradient, column wash, and re-equilibration steps.

Mass Spectrometry (MS) Conditions: Mass spec analysis was performed using a Thermo Scientific Orbirap Eclipse mass spectrometer with a variable window DIA method. The MS1 scans were acquired by Orbitrap at resolution of 60,000 and a mass range set to m/z 400-1400. The RF lens was set to 60%, and the AGC (Automatic Gain Control) target was normalized to a value of 200%. The absolute AGC value was set at 1.0e6, with a custom maximum injection time of 100 ms.

For the data independent MS/MS (tMS²) analysis, the MS2 scans were acquired in Orbitrap at the resolution of 15,000, the fragmentation of precursors was performed by using high-energy collisional dissociation (HCD) mode at the energy of 33%. The quadrupole was used for ion isolation, with isolation windows defined by a pre-determined table (m/z), the variable window definition is in the supplementary material (Table S1). The mass range for the MS/MS scans was set from m/z 200 to m/* 2000. The RF lens was maintained at 50%, and the AGC target was normalized to 1000%. The absolute AGC value was set at 5.00e5.

### HiBiT cellular degradation assay

MTH1-fusions (MTH1-RNF152, MTH1-LAPTM4a, MTH1-LAPTM5) stably expressing HCT116 (knock-in-HiBiT-FKBP12^F36V^-KRAS^G12D^ (HF-KRAS)) cells were seeded at a density of 10,000 cells per well in 96-well plates (Corning 3904). Next day, cells were treated with increasing concentrations of KRAS PROTAC, LYMTAC-2, LYMTAC-3, or LYMTAC-4. After 24 h, HiBiT assay was performed using Nano Glo-HiBit lytic reagent according to manufacturer’s protocol. The luminescence readings were recorded using EnVision plate reader and plotted as four parameter non-linear regression curve fit using GraphPad Prism 8.4.3.

### Mechanism of action studies using control compounds

HCT116 (HF-KRAS) cells stably expressing MTH1-fusion were seeded at a density of 5×10^5^ cells per well in 6-well dishes. Next day, cells were pre-treated with 200 nM BafA1, 2 µM BTZ or 10 µM MG-132, and 1 µM TAK-243 for 30 min and co-treated with DMSO or 500 nM LYMTAC-2 for 6 h. For PTK2 and EPHA2 rescue experiments, HEK-293T cells were pre-treated with 200 nM BafA1 and 10 µM MG-132 for 30 min and co-treated with DMSO or 500 nM LYMTAC-1 for 19 h. AsPC-1 cells were pre-treated with 200 nM BafA1 for 30 min, followed by 100 nM LYMTAC-4 co-treatment for 6 h. After treatment, cells were lysed in RIPA buffer and subjected to immunoblotting with indicated antibodies.

### Ligand competition experiments

Cells stably expressing MTH1-fusion were seeded at a density of 5×10^5^ cells per well in 6-well dishes. HEK293 cells stably expressing MTH1-RNF152 were pre-treated with 25 µM MTH1-ligand for 30 min followed by co-treatment with 500 nM LYMTAC-1 for 19 h. AsPC-1 cells were pretreated with 10 µM MTH1-ligand or 10 µM pan-KRAS inhibitor for 30 min, followed by 100 nM LYMTAC-4 co-treatment for 6 h. After treatment, cells were lysed and subjected to immunoblotting with indicated antibodies.

### Ternary complex formation assay

For ternary complex assays, AsPC-1 cells stably expressing FLAG-MTH1-RNF152 or FLAG-MTH1-LAPTM4a were seeded in 10-cm dishes at a density of 5×10^6^ cells. Next day, cells were pre-treated with BafA1 for 30 min, followed by co-treatment with DMSO, 1 µM LYMTAC-3, or 1 µM LYMTAC-4 for 4 h. Cells were harvested and lysed in IP-lysis buffer supplemented with protease inhibitor. Next, the cell lysate was subjected to immunoprecipitation with anti-FLAG magnetic beads overnight at 4 °C. Then, beads were washed three times with wash buffer (100 mM Tris–HCl pH 7.5, 150 mM NaCl, 0.1% Tween-20) and the samples were subjected to SDS-PAGE and Western blotting. The respective blots are probed with indicated antibodies.

### Ubiquitylation assays

To probe LYMTAC-induced KRAS ubiquitylation, HCT116 (HF-KRAS+MTH1 fusion) were seeded in a 10-cm dish at a cell density of 5×10^6^ cells. Next day, 4 µg HA-ubiquitin was transfected into HCT116 (HF-KRAS+MTH1 fusion) using Lipofectamine 2000. After 24 h, cells were pre-treated with 200 nM BafA1 for 30min followed by co-treatment with DMSO or 1 µM LYMTAC-2 for 4 h. Cells were then placed on ice, washed with ice-cold 1X PBS, and lysed in IP lysis buffer supplemented with 1X protease inhibitor cocktail, 5 mM 1,10-phenanthroline monohydrate, 10 mM N-ethylmaleimide, and 20 µM PR-619. Then, the lysate was incubated with 2 µg of anti-HiBiT antibody for 20 min at room temperature. HiBiT antibody incubated lysate was next added to 20 µL of protein G Sepharose magnetic beads and incubated overnight. After incubation, beads were washed 3X with wash buffer and bound proteins were eluted using 2X NuPAGE™ LDS Sample Buffer supplemented with 10% β-ME. Proteins were separated in SDS-PAGE and immunoblotted with respective antibodies.

### Confocal microscopy

For microscopy experiments, mNeonGreen-KRAS^WT^ cells stably expressing FLAG-MTH1-RNF152 were seeded at a density of 15,000 cells per well in 8-well chambered dishes (Ibidi). Next day, cells were pre-treated with 200 nM BafA1 for 30 min and co-treated with 1 µM LYMTAC-4 for 4 h. Then, cells were washed 2X with PBS and fixed with 4% paraformaldehyde in PBS for 15 min at room temperature. After fixing, cells were permeabilized with 0.2% Triton-X-100 in PBS for 10 min at room temperature. Then, cells were incubated with blocking buffer (3% BSA, 0.2% Triton-X-100 in PBS) for 1 h at room temperature followed by overnight incubation at 4 °C with FLAG (1:300 dilution) and LAMP1 (1:50 dilution) antibodies in antibody dilution buffer (1% BSA, 0.2% Triton-X-100 in PBS). Next day, cells were washed 3X with PBS at room temperature and incubated with secondary antibodies (1:500 dilution, Alexa Fluor 568 and Alexa Fluro 647) at room temperature for 1 h. Next, cells were washed 3X with PBS and stained with Hoechst stain (3 μg/mL) in PBS for 10 min at room temperature followed by two PBS washes. The cells were imaged using a 63x/1.30 NA glycerol immersion objective lens on an inverted Leica TCS SP8 confocal microscope and images were analyzed using Leica LAS X software.

### Washout experiment with LYMTAC-4 in AsPC-1

AsPC-1 cells stably expressing RNF152 were seeded at a density of 4×10^5^ cells per well in 6-well dishes. After 24 h, cells were treated with DMSO, 100 nM pan-KRAS inhibitor, or 100 nM LYMTAC-4 for 4 h, removed media, washed three times with PBS, and replaced with fresh media for 48 h. For LYMTAC-4 treated wells, fresh media was added in the absence or presence of 5 μM MTH1-ligand for 48 h. After 48 h, cells were lysed and subjected to SDS-PAGE and immunoblotting with respective antibodies.

### Cell proliferation assays

AsPC-1 cells stably expressing RNF152 were seeded at a density of 2000 cells per well in 96-well plates. Next day, cells were treated with dose titrations of pan-KRASi, KRAS PROTAC, or LYMTAC-4. After five days of treatment, cell viability was measured using CyQUANT™ direct cell proliferation assay reagents following manufacturer’s protocol. The fluorescence readings were recorded using EnVision plate reader and plotted as four parameter non-linear regression curve fit using GraphPad Prism 8.4.3.

## Supplementary Figures

**Figure S1:**
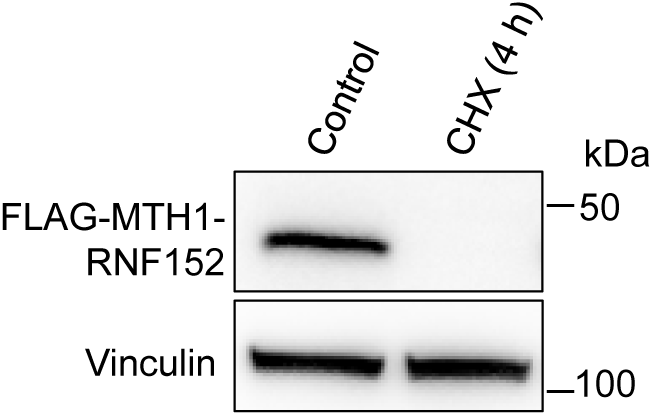
MTH1-RNF152 is a short-lived LMP in HEK293T cells. Cycloheximide (CHX) assay in MTH1-RNF152 stably expressing HEK293T cells. Cells were treated with 100 µg/ml CHX for 4 h. Cells were harvested at 0 h to include as a control. Cells were lysed and subjected to immunoblotting with MTH1 and vinculin antibodies. Data are representative of two independent experiments.

**Figure S2:**
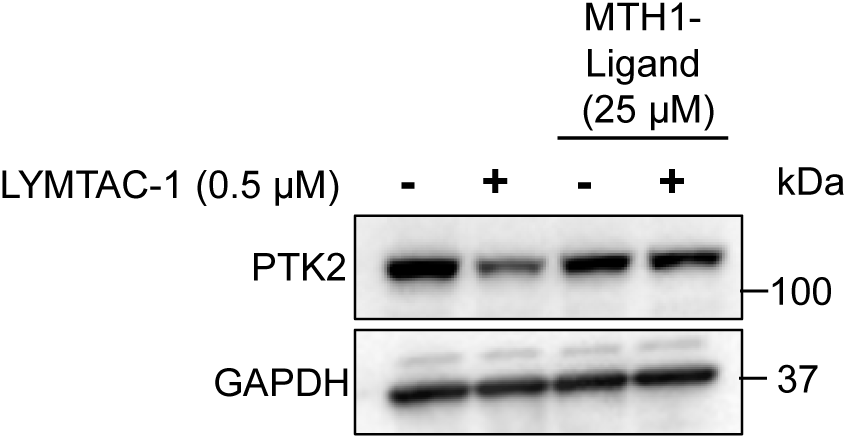
LYMTAC-1 induced PTK2 degradation is RNF152-dependent. HEK293T cells stably expressing FLAG-MTH1-RNF152 were pre-treated with 25 µM MTH1 ligand for 30 min, followed by DMSO or 500 nM LYMTAC-1 treatment for 19 h. Cells were lysed and subjected to immunoblotting with indicated antibodies. Data are representative of two independent experiments.

**Figure S3:**
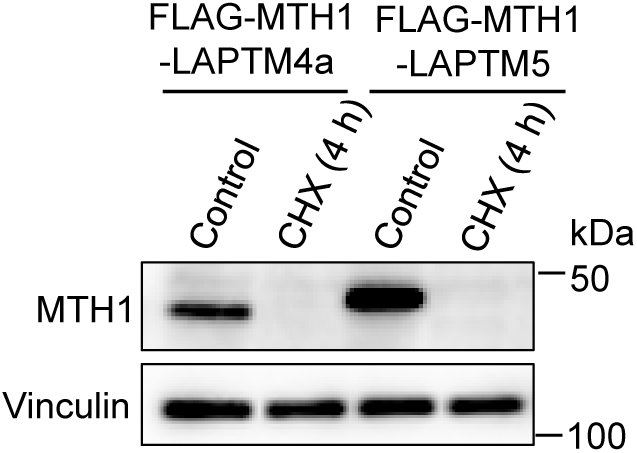
MTH1-LAPTM4a and MTH1-LAPTM5 are short-lived LMPs in HEK293T cells. Cells were treated with 100 µg/ml CHX for 4 h. Cells were harvested at 0 h to include as a control. Cells were lysed and subjected to immunoblotting with MTH1 and vinculin antibodies. Data are representative of two independent experiments.

**Figure S4:**
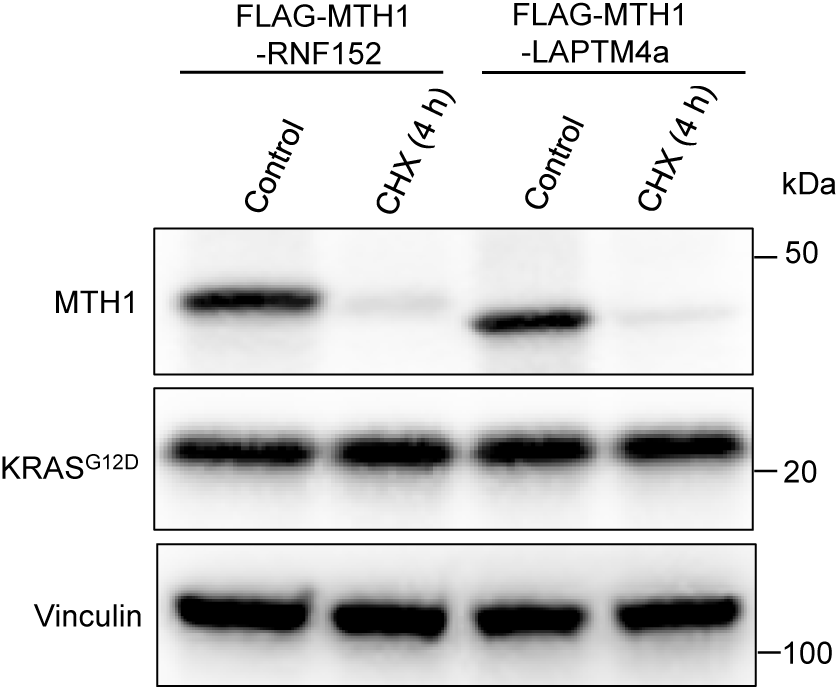
MTH1-RNF152 and MTH1-LAPTM4a are short-lived LMPs in AsPC-1 cells. AsPC-1 cells were treated with 100 µg/ml CHX for 4 h. Cells were harvested at 0 h to include as a control. Cells were lysed and subjected to immunoblotting with MTH1, KRAS, and vinculin antibodies. Data are representative of two independent experiments.

**Figure S5:**
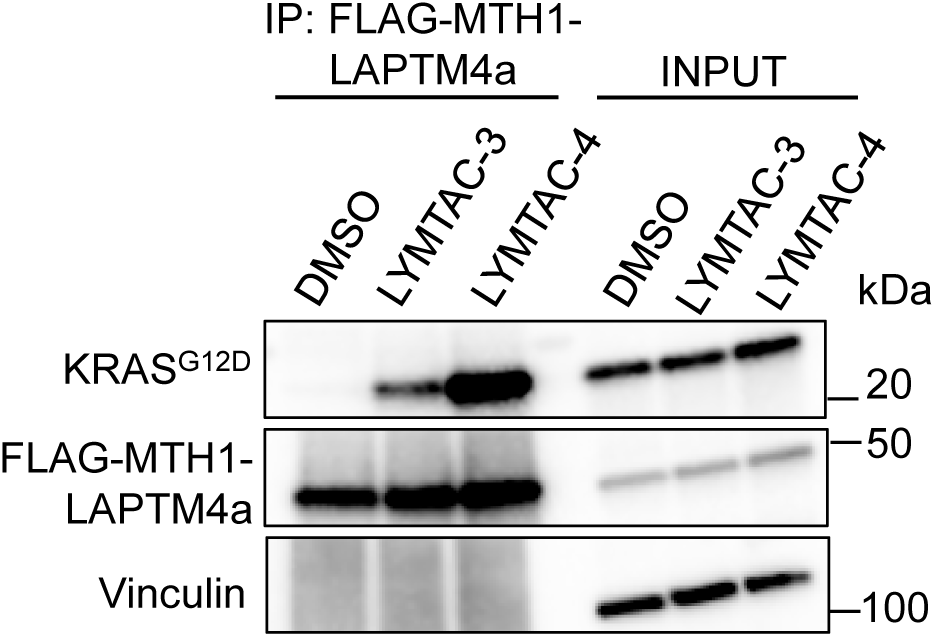
LYMTAC-4 induces a stable ternary complex between KRAS^G12D^ and LAPTM4a in AsPC-1 cells. AsPC-1 cells stably expressing FLAG-MTH1-LAPTM4a were pre-treated with Bafilomycin A1 for 30 min, followed by co-treatment with DMSO, 1 µM LYMTAC-3, or LYMTAC-4 for 4 h and subjected to immunoprecipitation with anti-FLAG antibody. The respective blots are probed with KRAS, MTH1, and vinculin antibodies. Data are representative of two independent experiments. Whole cell lysate was used as INPUT.

**Figure S6:**
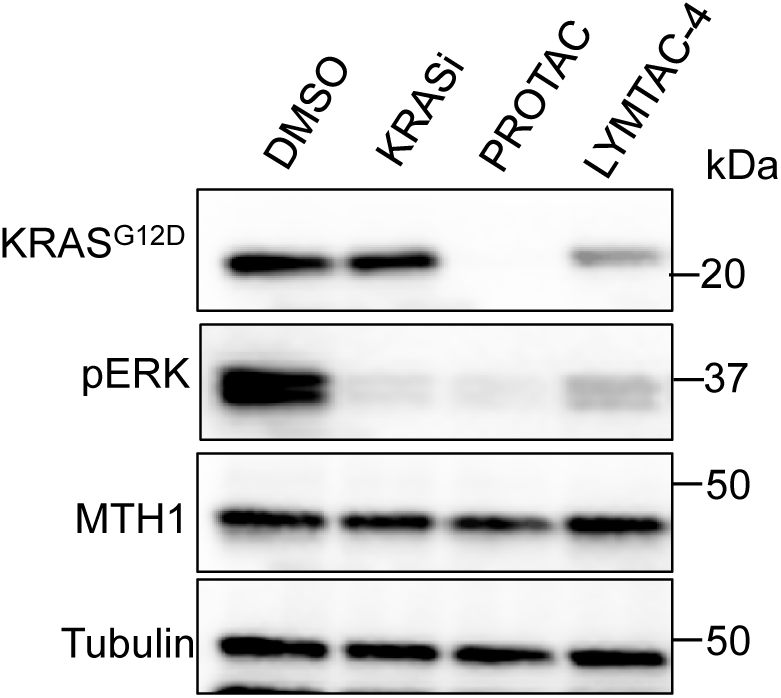
LAPTM4a-recruiting LYMTAC-4 induces KRAS^G12D^ degradation and suppresses p-ERK signaling in AsPC-1 cells. AsPC-1 cells stably expressing MTH1-LAPTM4a were treated with DMSO or 100 nM of the indicated compounds for 24 h and subjected to immunoblotting with KRAS, pERK, MTH1, and tubulin antibodies. Data are representative of two independent experiments.

**Table S1:** Mass List Table.

| Adduct | m/z | z | t start (min) | t stop (min) | Isolation Window (m/z) |
| --- | --- | --- | --- | --- | --- |
| (no adduct) | 404.85 | 3 | 0 | 104 | 10.7 |
| (no adduct) | 413.95 | 3 | 0 | 104 | 9.5 |
| (no adduct) | 421.95 | 3 | 0 | 104 | 8.5 |
| (no adduct) | 429.2 | 3 | 0 | 104 | 8 |
| (no adduct) | 436.15 | 3 | 0 | 104 | 7.9 |
| (no adduct) | 442.9 | 3 | 0 | 104 | 7.6 |
| (no adduct) | 449.25 | 3 | 0 | 104 | 7.1 |
| (no adduct) | 455.55 | 3 | 0 | 104 | 7.5 |
| (no adduct) | 462.05 | 3 | 0 | 104 | 7.5 |
| (no adduct) | 468.3 | 3 | 0 | 104 | 7 |
| (no adduct) | 474.45 | 3 | 0 | 104 | 7.3 |
| (no adduct) | 480.4 | 3 | 0 | 104 | 6.6 |
| (no adduct) | 486.25 | 3 | 0 | 104 | 7.1 |
| (no adduct) | 492.5 | 3 | 0 | 104 | 7.4 |
| (no adduct) | 498.5 | 3 | 0 | 104 | 6.6 |
| (no adduct) | 503.55 | 3 | 0 | 104 | 5.5 |
| (no adduct) | 508.3 | 3 | 0 | 104 | 6 |
| (no adduct) | 513.3 | 3 | 0 | 104 | 6 |
| (no adduct) | 518.3 | 3 | 0 | 104 | 6 |
| (no adduct) | 523.2 | 3 | 0 | 104 | 5.8 |
| (no adduct) | 528.2 | 3 | 0 | 104 | 6.2 |
| (no adduct) | 533.4 | 3 | 0 | 104 | 6.2 |
| (no adduct) | 538.65 | 3 | 0 | 104 | 6.3 |
| (no adduct) | 543.8 | 3 | 0 | 104 | 6 |
| (no adduct) | 548.8 | 3 | 0 | 104 | 6 |
| (no adduct) | 553.8 | 3 | 0 | 104 | 6 |
| (no adduct) | 559.05 | 3 | 0 | 104 | 6.5 |
| (no adduct) | 564.55 | 3 | 0 | 104 | 6.5 |
| (no adduct) | 570.05 | 3 | 0 | 104 | 6.5 |
| (no adduct) | 575.35 | 3 | 0 | 104 | 6.1 |
| (no adduct) | 580.65 | 3 | 0 | 104 | 6.5 |
| (no adduct) | 586.1 | 3 | 0 | 104 | 6.4 |
| (no adduct) | 591.8 | 3 | 0 | 104 | 7 |
| (no adduct) | 597.8 | 3 | 0 | 104 | 7 |
| (no adduct) | 603.3 | 3 | 0 | 104 | 6 |
| (no adduct) | 608.7 | 3 | 0 | 104 | 6.8 |
| (no adduct) | 614.75 | 3 | 0 | 104 | 7.3 |
| (no adduct) | 621.1 | 3 | 0 | 104 | 7.4 |
| (no adduct) | 627.3 | 3 | 0 | 104 | 7 |
| (no adduct) | 633.55 | 3 | 0 | 104 | 7.5 |
| (no adduct) | 640.25 | 3 | 0 | 104 | 7.9 |
| (no adduct) | 647 | 3 | 0 | 104 | 7.6 |
| (no adduct) | 653.8 | 3 | 0 | 104 | 8 |
| (no adduct) | 660.8 | 3 | 0 | 104 | 8 |
| (no adduct) | 667.8 | 3 | 0 | 104 | 8 |
| (no adduct) | 675.05 | 3 | 0 | 104 | 8.5 |
| (no adduct) | 682.6 | 3 | 0 | 104 | 8.6 |
| (no adduct) | 690.15 | 3 | 0 | 104 | 8.5 |
| (no adduct) | 697.4 | 3 | 0 | 104 | 8 |
| (no adduct) | 704.85 | 3 | 0 | 104 | 8.9 |
| (no adduct) | 713.1 | 3 | 0 | 104 | 9.6 |
| (no adduct) | 721.9 | 3 | 0 | 104 | 10 |
| (no adduct) | 730.9 | 3 | 0 | 104 | 10 |
| (no adduct) | 740.1 | 3 | 0 | 104 | 10.4 |
| (no adduct) | 749.6 | 3 | 0 | 104 | 10.6 |
| (no adduct) | 758.9 | 3 | 0 | 104 | 10 |
| (no adduct) | 768.4 | 3 | 0 | 104 | 11 |
| (no adduct) | 777.9 | 3 | 0 | 104 | 10 |
| (no adduct) | 787.1 | 3 | 0 | 104 | 10.4 |
| (no adduct) | 796.6 | 3 | 0 | 104 | 10.6 |
| (no adduct) | 806.4 | 3 | 0 | 104 | 11 |
| (no adduct) | 816.7 | 3 | 0 | 104 | 11.6 |
| (no adduct) | 827.25 | 3 | 0 | 104 | 11.5 |
| (no adduct) | 838.45 | 3 | 0 | 104 | 12.9 |
| (no adduct) | 850.15 | 3 | 0 | 104 | 12.5 |
| (no adduct) | 861.7 | 3 | 0 | 104 | 12.6 |
| (no adduct) | 873.45 | 3 | 0 | 104 | 12.9 |
| (no adduct) | 885.9 | 3 | 0 | 104 | 14 |
| (no adduct) | 898.1 | 3 | 0 | 104 | 12.4 |
| (no adduct) | 910.15 | 3 | 0 | 104 | 13.7 |
| (no adduct) | 923.5 | 3 | 0 | 104 | 15 |
| (no adduct) | 937.45 | 3 | 0 | 104 | 14.9 |
| (no adduct) | 951.45 | 3 | 0 | 104 | 15.1 |
| (no adduct) | 965.75 | 3 | 0 | 104 | 15.5 |
| (no adduct) | 980.7 | 3 | 0 | 104 | 16.4 |
| (no adduct) | 995.8 | 3 | 0 | 104 | 15.8 |
| (no adduct) | 1011.85 | 3 | 0 | 104 | 18.3 |
| (no adduct) | 1029 | 3 | 0 | 104 | 18 |
| (no adduct) | 1046.8 | 3 | 0 | 104 | 19.6 |
| (no adduct) | 1065.8 | 3 | 0 | 104 | 20.4 |
| (no adduct) | 1086.3 | 3 | 0 | 104 | 22.6 |
| (no adduct) | 1108.25 | 3 | 0 | 104 | 23.3 |
| (no adduct) | 1131.95 | 3 | 0 | 104 | 26.1 |
| (no adduct) | 1157.05 | 3 | 0 | 104 | 26.1 |
| (no adduct) | 1184.3 | 3 | 0 | 104 | 30.4 |
| (no adduct) | 1212.25 | 3 | 0 | 104 | 27.5 |
| (no adduct) | 1242.1 | 3 | 0 | 104 | 34.2 |
| (no adduct) | 1278.2 | 3 | 0 | 104 | 40 |
| (no adduct) | 1320.1 | 3 | 0 | 104 | 45.8 |
| (no adduct) | 1371.25 | 3 | 0 | 104 | 58.5 |

## Supporting Information (Chemical Synthesis)

### LYMTAC-1

**6-(6-((2-(2-(2-(2-(4-(4-((5-chloro-4-((2-(isopropyl sulfonyl) phenyl) amino) pyrimidin-2-yl) amino) phenyl) piperazin-1-yl) ethoxy) ethoxy) ethoxy) ethyl) carbamoyl) pyridin-3-yl)-N-methyl-4-(phenylamino) quinoline-3-carboxamide**

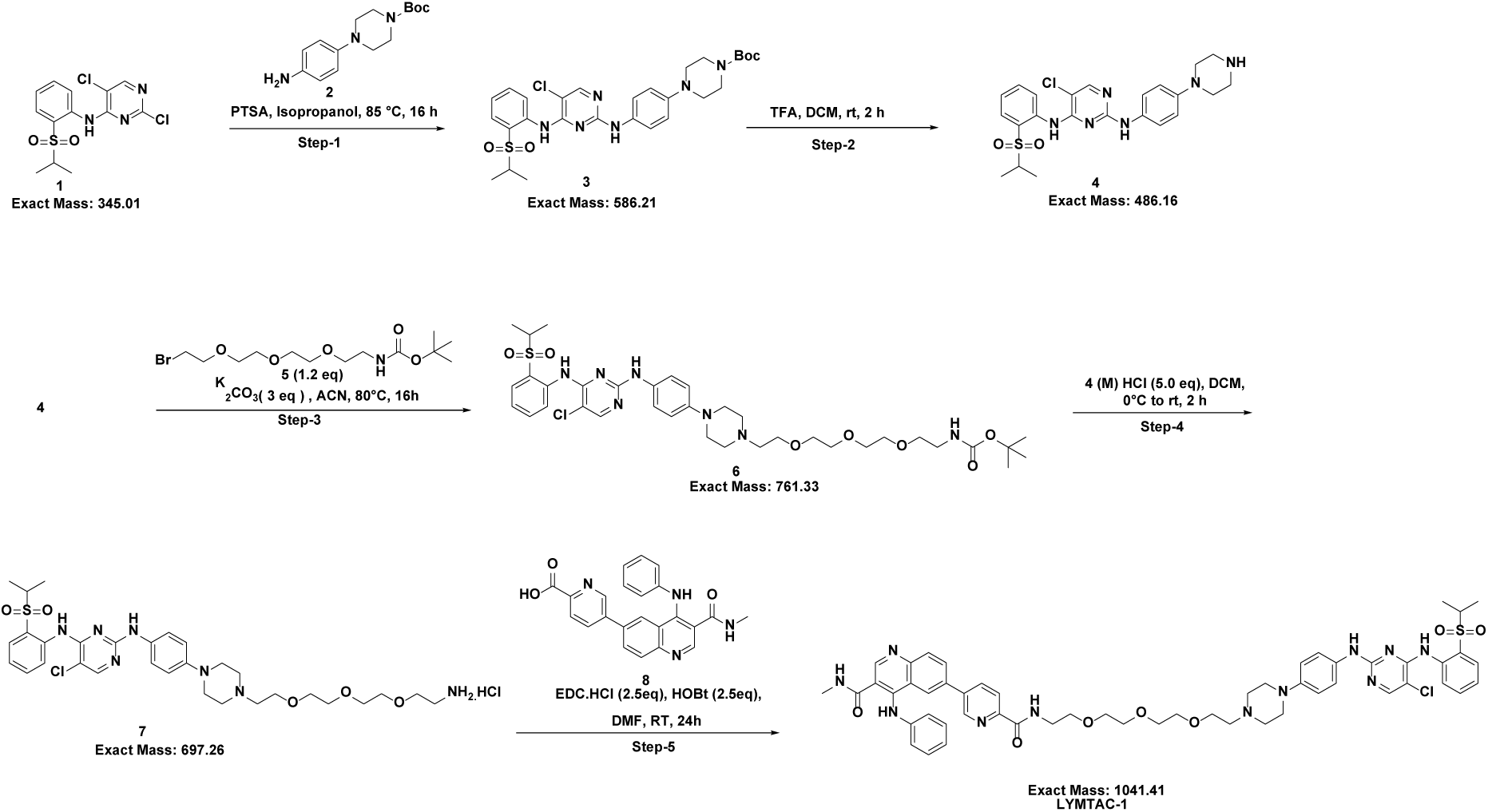

**Step-1: Synthesis of tert-butyl 4-(4-((5-chloro-4-((2-(isopropyl sulfonyl) phenyl) amino) pyrimidin-2-yl) amino) phenyl) piperazine-1-carboxylate**

To a 100-mL round-bottomed flask was added 2,5-dichloro-N-(2-(isopropyl sulfonyl) phenyl) pyrimidin-4-amine (2 g, 5.78 mmol, 1.0 eq) and tert-butyl 4-(4-aminophenyl) piperazine-1-carboxylate (1.602 g, 5.78 mmol, 1.0 eq) followed by addition of *p*-toluene sulfonic acid monohydrate (1.319 g, 6.93 mmol, 1.2 eq) in isopropanol (40.0 mL) to the reaction mixture at room temperature and stirred at 100°C for 16 h. After completion, the reaction mixture was concentrated in vacuo to give the crude tert-butyl 4-(4-((5-chloro-4-((2-(isopropyl sulfonyl) phenyl) amino) pyrimidin-2-yl) amino) phenyl) piperazine-1-carboxylate as a light-yellow oil.

ESI [M-100]: 487.2.

**Step-2: Synthesis of 5-chloro-N4-(2-(isopropyl sulfonyl) phenyl)-N2-(4-(piperazin-1-yl) phenyl) pyrimidine-2,4-diamine**

To a 100-mL round-bottomed flask was added crude tert-butyl 4-(4-((5-chloro-4-((2-(isopropyl sulfonyl) phenyl) amino) pyrimidin-2-yl) amino) phenyl) piperazine-1-carboxylate (3 g, 2.55 mmol) in dichloromethane (60 mL), followed by addition of trifluoroacetic acid (0.197 mL, 2.55 mmol, 1.0 eq) at 25°C. The reaction mixture was stirred at room temperature for 1 h. After completion, the reaction mixture was concentrated and quenched with water (50 mL), extracted with dichloromethane (2 x 50 mL). The aqueous layer was basified with saturated NaOH solution and extracted with ethyl acetate (2 x 50 mL). The organic layer was washed with brine and dried over Na2SO4, filtered and concentrated under reduced pressure to provide 5-chloro-N4-(2-(isopropyl sulfonyl) phenyl)-N2-(4-(piperazin-1-yl) phenyl) pyrimidine-2,4-diamine as a blue solid.

***m/z* (ESI):** 487.1 (M+1) ^+^

ESI [M+H]^+^: 487.1

**Step-3: Synthesis of tert-butyl (2-(2-(2-(2-(4-(4-((5-chloro-4-((2-(isopropyl sulfonyl) phenyl) amino) pyrimidin-2-yl) amino) phenyl) piperazin-1-yl) ethoxy) ethoxy) ethoxy) ethyl) carbamate**

To a 50 mL round-bottomed flask was added 5-chloro-N4-(2-(isopropyl sulfonyl) phenyl)-N2-(4-(piperazin-1-yl) phenyl) pyrimidine-2,4-diamine (500 mg, 1.027 mmol, 1.0 eq), tert-butyl (2-(2-(2-(2-bromoethoxy) ethoxy) ethoxy) ethyl) carbamate (439 mg, 1.232 mmol, 1.2 eq) and K2CO3 (426 mg, 3.08 mmol, 3.0 eq) in acetonitrile (10 mL) and stirred at 80 °C for 16h. After completion, the reaction mass was filtered through celite bed, washed with DCM and concentrated under reduced pressure. The crude material was absorbed onto a plug of silica gel and purified by chromatography through a silica gel column, eluting with a gradient of 0% to 12% MeOH in CH2Cl2 to provide tert-butyl (2-(2-(2-(2-(4-(4-((5-chloro-4-((2-(isopropyl sulfonyl) phenyl) amino) pyrimidin-2-yl) amino) phenyl) piperazin-1-yl) ethoxy) ethoxy) ethoxy) ethyl) carbamate (500 mg, 0.656 mmol, 63.9 % yield) as a grey sticky solid.

^1^H NMR (400 MHz, DMSO-*d*6) δ 9.49 (s, 1H), 9.31 (s, 1H), 8.66 (s, 1H), 8.24 (s, 1H), 7.84 (dd, J = 7.9, 1.6 Hz, 1H), 7.74 (t, J = 7.8 Hz, 1H), 7.43 (d, J = 8.3 Hz, 2H), 7.40 – 7.35 (m, 1H), 6.87 (d, J = 8.4 Hz, 2H), 6.76 (s, 1H), 3.74 (t, J = 5.8 Hz, 1H), 3.60 – 3.48 (m, 14H), 3.42 – 3.36 (m, 5H), 1.37 (s, 9H), 1.17 (d, J = 6.8 Hz, 6H). ESI [M+H]^+^: 762.3

**Step-4: Synthesis of N2-(4-(4-(2-(2-(2-(2-aminoethoxy) ethoxy) ethoxy) ethyl) piperazin-1-yl) phenyl)-5-chloro-N4-(2-(isopropyl sulfonyl) phenyl) pyrimidine-2,4-diamine hydrochloride**

To a 25 mL round-bottomed flask was added tert-butyl (2-(2-(2-(2-(4-(4-((5-chloro-4-((2-(isopropyl sulfonyl) phenyl) amino) pyrimidin-2-yl) amino) phenyl) piperazin-1-yl) ethoxy) ethoxy) ethoxy) ethyl) carbamate (500 mg, 0.656 mmol, 1.0 eq) in DCM (5 mL) followed by addition of 4 M HCl in Dioxane (820 µL, 3.28 mmol, 5.0 eq) to the reaction mixture at 0°C and stirred at room temperature for 2 h. After completion, the reaction mixture was concentrated under reduced pressure followed by trituration with di-ethyl ether to afford 500 mg of crude N2-(4-(4-(2-(2-(2-(2-aminoethoxy) ethoxy) ethoxy) ethyl) piperazin-1-yl) phenyl)-5-chloro-N4-(2-(isopropyl sulfonyl) phenyl) pyrimidine-2,4-diamine hydrochloride as a blackish solid which was directly used in the next step. ^1^H NMR (400 MHz, DMSO-*d*6): δ 10.74 (s, 1H), 9.57 (s, 1H), 9.49 (s, 1H), 7.95 (s, 2H), 7.51 – 7.36 (m, 1H), 6.94 (d, J = 9.0 Hz, 1H), 4.16 (s, 11H), 3.88 (t, J = 5.0 Hz, 1H), 3.77 – 3.69 (m, 1H), 3.68 – 3.56 (m, 9H), 3.56 – 3.40 (m, 1H), 3.37 (d, J = 5.7 Hz, 1H), 3.17 (dt, J = 41.9, 11.7 Hz, 2H), 2.97 (q, J = 5.6 Hz, 1H), 1.17 (d, J = 6.8 Hz, 2H). ESI [M+H]^+^: 663.1

**Step-5: Synthesis of 6-(6-((2-(2-(2-(2-(4-(4-((5-chloro-4-((2-(isopropyl sulfonyl) phenyl) amino) pyrimidin-2-yl) amino) phenyl) piperazin-1-yl) ethoxy) ethoxy) ethoxy) ethyl) carbamoyl) pyridin-3-yl)-N-methyl-4-(phenylamino) quinoline-3-carboxamide (LYMTAC-1)**

To a 25-mL round-bottomed flask was added 5-(3-(methylcarbamoyl)-4-(phenylamino)quinolin-6-yl)picolinic acid (60 mg, 0.151 mmol, 1.0 eq), N2-(4-(4-(2-(2-(2-(2-aminoethoxy) ethoxy) ethoxy) ethyl) piperazin-1-yl) phenyl)-5-chloro-N4-(2-(isopropyl sulfonyl) phenyl) pyrimidine-2,4-diamine hydrochloride (126 mg, 0.181 mmol, 1.2 eq), EDC (72.2 mg, 0.376 mmol, 2.5 eq) and HOBt (57.7 mg, 0.376 mmol, 2.5 eq) in dichloromethane (3.0 mL) followed by addition of DIPEA (105 µL, 0.602 mmol, 4.0 eq) to the reaction mixture and stirred at room temperature for 12 h. After completion, the reaction mixture was diluted with water (5 mL) and extracted with 10 % Methanol in CH2Cl2 (3 x 30 mL). The organic extract was washed with brine (1x 5mL) and dried over Na2SO4, filtered and concentrated in vacuo to give the crude material as a light-yellow oil which was purified by prep-HPLC using Discovery C-18 (250*21.2) mm 5.0 µm column with a mobile phase of A: 0.1% Formic acid water B: ACN using a flow rate of 15 mL/min, followed by lyophilization to provide 6-(6-((2-(2-(2-(2-(4-(4-((5-chloro-4-((2-(isopropylsulfonyl)phenyl)amino)pyrimidin-2-yl)amino)phenyl)piperazin-1-yl)ethoxy)ethoxy)ethoxy)ethyl)carbamoyl)pyridin-3-yl)-N-methyl-4-(phenylamino)quinoline-3-carboxamide (61.06 mg, 0.059 mmol, 38.9 % yield) as a light yellow solid.

^1^H NMR (400 MHz, DMSO-*d*6): δ 10.09 (s, 1H), 9.48 (s, 1H), 9.29 (s, 1H), 8.83 (s, 1H), 8.80 – 8.68 (m, 2H), 8.64 (s, 1H), 8.52 (q, J = 4.5 Hz, 1H), 8.36 (d, J = 2.0 Hz, 1H), 8.24 – 8.12 (m, 3H), 8.07 (dd, J = 13.5, 8.5 Hz, 2H), 7.83 (dd, J = 8.0, 1.6 Hz, 1H), 7.77 – 7.68 (m, 1H), 7.43 – 7.27 (m, 5H), 7.07 (dd, J = 8.3, 7.1 Hz, 3H), 6.85 – 6.78 (m, 2H), 3.62 – 3.38 (m, 17H), 3.01 (t, J = 5.0 Hz, 4H), 2.57 (d, J = 4.5 Hz, 3H), 2.47 (d, J = 6.0 Hz, 1H), 1.16 (d, J = 6.8 Hz, 6H). ESI [M+H]^+^: 1042.2

### LYMTAC-2

**(R)-3-(3,4-dimethoxyphenyl)-1-(3-((1-(5-(3-(methylcarbamoyl)-4-(phenylamino)quinolin-6-yl)pyridin-2-yl)-1,13-dioxo-5,8,11-trioxa-2,14-diazahexadecan-16-yl)oxy)phenyl)propyl (S)-1-((S)-2-(3,4,5-trimethoxyphenyl)butanoyl)piperidine-2-carboxylate**

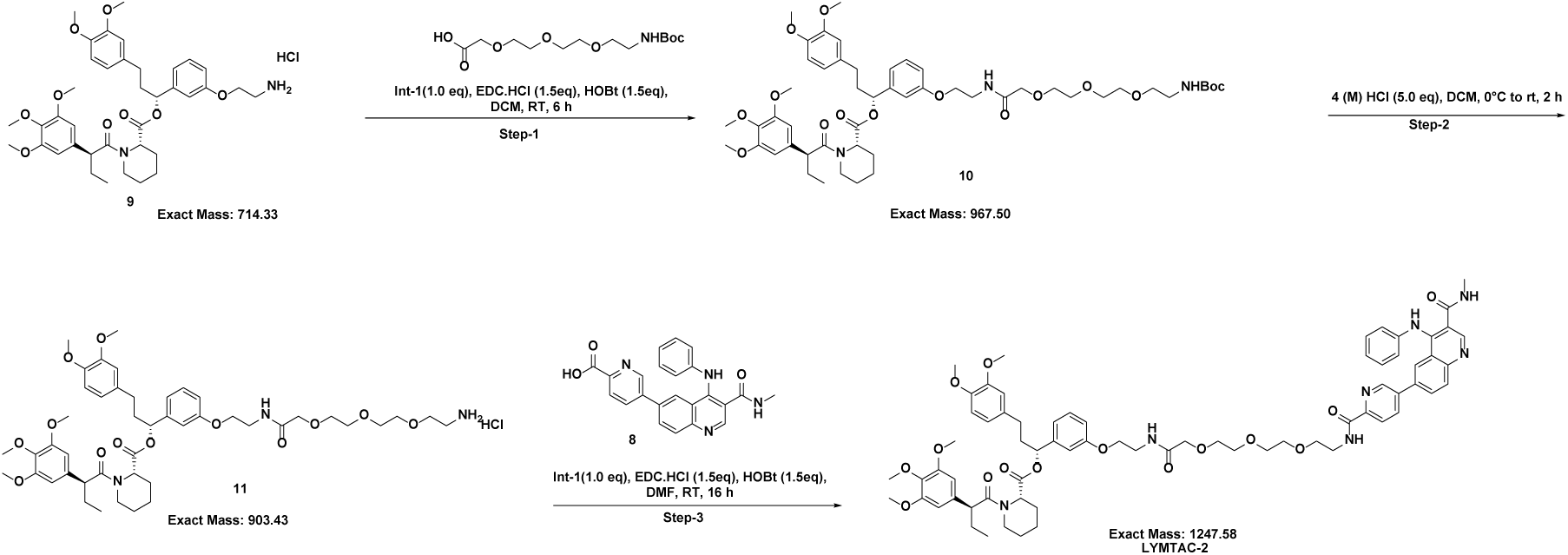

**Step-1: Synthesis of (R)-3-(3,4-dimethoxyphenyl)-1-(3-((2,2-dimethyl-4,16-dioxo-3,8,11,14-tetraoxa-5,17-diazanonadecan-19-yl)oxy)phenyl)propyl(S)-1-((S)-2-(3,4,5-trimethoxyphenyl)butanoyl)piperidine-2-carboxylate**

To a solution of (R)-1-(3-(2-aminoethoxy)phenyl)-3-(3,4-dimethoxyphenyl)propyl (S)-1-((S)-2-(3,4,5-trimethoxyphenyl)butanoyl)piperidine-2-carboxylate hydrochloride (200 mg, 0.280 mmol) and 2,2-dimethyl-4-oxo-3,8,11,14-tetraoxa-5-azahexadecan-16-oic acid (95 mg, 0.308 mmol) in dichloromethane (30 mL) was added 1H-benzo[d][1,2,3]triazol-1-ol (56.7 mg, 0.419 mmol), EDC (80 mg, 0.419 mmol) and DIPEA (293 µL, 1.678 mmol) at RT and stirred for 6 h. The reaction mixture was diluted with water (10 mL) and extracted with DCM (2 x 20 mL). The organic extract was washed with water (15 mL) and dried over Na2SO4. The solution was filtered and concentrated to give the crude material as an orange oil. The crude material was absorbed onto a plug of silica gel and purified by column chromatography through a Redi-Sep pre-packed silica gel column (40 g), eluting with a gradient of 10%to 15% MeOH in CH2Cl2, to provide (R)-3-(3,4-dimethoxyphenyl)-1-(3-((2,2-dimethyl-4,16-dioxo-3,8,11,14-tetraoxa-5,17-diazanonadecan-19-yl)oxy)phenyl)propyl (S)-1-((S)-2-(3,4,5-trimethoxyphenyl)butanoyl)piperidine-2-carboxylate (190 mg, 0.196 mmol, 70 % yield) as colorless thick oil. ^1^H NMR (400 MHz, DMSO-*d*6): δ 7.85 (t, J = 6.1 Hz, 1H), 7.31 (t, J = 7.9 Hz, 1H), 7.16 (t, J = 7.9 Hz, 1H), 6.93 (d, J = 13.5 Hz, 1H), 6.88 – 6.82 (m, 2H), 6.79 (d, J = 1.9 Hz, 1H), 6.77 – 6.71 (m, 2H), 6.67 – 6.60 (m, 1H), 6.55 (d, J = 13.5 Hz, 2H), 5.77 (d, J = 7.4 Hz, 1H), 5.53 (dd, J = 8.3, 5.2 Hz, 1H), 5.28 (d, J = 5.3 Hz, 1H), 5.05 (s, 1H), 4.41 (s, 1H), 4.09 (q, J = 5.3 Hz, 1H), 4.01 (t, J = 5.8 Hz, 2H), 3.91 (s, 2H), 3.86 (t, J = 7.1 Hz, 1H), 3.75 (s, 1H), 3.71 (d, J = 3.3 Hz, 5H), 3.64 (s, 1H), 3.61 – 3.54 (m, 9H), 3.50 (q, J = 5.9, 4.5 Hz, 5H), 3.17 (d, J = 5.3 Hz, 1H), 3.05 (q, J = 6.0 Hz, 2H), 2.63 (d, J = 12.4 Hz, 1H), 2.39 (t, J = 7.8 Hz, 2H), 2.17 (d, J = 13.0 Hz, 1H), 1.93 (dq, J = 21.0, 8.1, 7.5 Hz, 3H), 1.63 (s, 2H), 1.56 (dt, J = 14.8, 7.7 Hz, 2H), 1.36 (s, 10H), 1.25 (d, J = 8.5 Hz, 2H), 1.13 (d, J = 13.5 Hz, 1H), 0.81 (t, J = 7.3 Hz, 2H). ESI [M+H]^+^: 968.4

**Step-2: Synthesis of (R)-1-(3-((14-amino-4-oxo-6,9,12-trioxa-3-azatetradecyl)oxy)phenyl)-3-(3,4-dimethoxyphenyl)propyl(S)-1-((S)-2-(3,4,5-trimethoxyphenyl)butanoyl)piperidine-2-carboxylate hydrochloride**

To a solution of (R)-3-(3,4-dimethoxyphenyl)-1-(3-((2,2-dimethyl-4,16-dioxo-3,8,11,14-tetraoxa-5,17-diazanonadecan-19-yl)oxy)phenyl)propyl (S)-1-((S)-2-(3,4,5-trimethoxyphenyl)butanoyl)piperidine-2-carboxylate (190 mg, 0.196 mmol) in dichloromethane (7.4 mL) was added HCl (4 M in 1,4-Dioxane) (5.96 µL, 0.196 mmol) at 0 °C . The reaction mixture was stirred at RT for 2 h and was concentrated in vacuo to give the crude material as an orange oil. The crude material was washed with MTBE to provide (R)-1-(3-((14-amino-4-oxo-6,9,12-trioxa-3-azatetradecyl)oxy)phenyl)-3-(3,4-dimethoxyphenyl)propyl (S)-1-((S)-2-(3,4,5-trimethoxyphenyl)butanoyl)piperidine-2-carboxylate hydrochloride (146 mg, 0.161 mmol, 82 % yield) as a light-yellow waxy solid. ESI [M+H]^+^: 868.3

**Step-3: Synthesis of (R)-3-(3,4-dimethoxyphenyl)-1-(3-((1-(5-(3-(methylcarbamoyl)-4-(phenylamino)quinolin-6-yl)pyridin-2-yl)-1,13-dioxo-5,8,11-trioxa-2,14-diazahexadecan-16-yl)oxy)phenyl)propyl (S)-1-((S)-2-(3,4,5-trimethoxyphenyl)butanoyl)piperidine-2-carboxylate (LYMTAC-2)**

To a solution of 5-(3-(methylcarbamoyl)-4-(phenylamino)quinolin-6-yl)picolinic acid (60 mg, 0.151 mmol) and (R)-1-(3-((14-amino-4-oxo-6,9,12-trioxa-3-azatetradecyl)oxy)phenyl)-3-(3,4-dimethoxyphenyl)propyl (S)-1-((S)-2-(3,4,5-trimethoxyphenyl)butanoyl)piperidine-2-carboxylate hydrochloride (136 mg, 0.151 mmol) in N, N-dimethylformamide (30 mL) was added 1H-benzo[d][1,2,3]triazol-1-ol (30.5 mg, 0.226 mmol), EDC (43.3 mg, 0.226 mmol) and DIPEA (158 µL, 0.904 mmol) at RT and stirred at RT for 6 h. The reaction mixture was diluted with water (10 mL) and extracted with CH2Cl2 (2 x 20 mL). The organic extract was washed with water (15 mL) and dried over Na_2_SO_4_. The solution was filtered and concentrated in vacuo to give the crude material as an orange oil. The crude material was purified by reverse-phase preparative HPLC using a kinetex C8 (250*21.2) mm 5.0µm, 0.1% FA in water B: ACN to provide (R)-3-(3,4-dimethoxyphenyl)-1-(3-((1-(5-(3-(methylcarbamoyl)-4-(phenylamino)quinolin-6-yl)pyridin-2-yl)-1,13-dioxo-5,8,11-trioxa-2,14-diazahexadecan-16-yl)oxy)phenyl)propyl (S)-1-((S)-2-(3,4,5-trimethoxyphenyl)butanoyl)piperidine-2-carboxylate (45 mg, 0.036 mmol, 23.94 % yield) as a yellow solid. ^d^ δ 10.07 (s, 1H), 8.83 (s, 1H),

8.77 (d, *J* = 2.3 Hz, 1H), 8.71 (t, *J* = 5.8 Hz, 1H), 8.52 (d, *J* = 4.5 Hz, 1H), 8.36 (d, *J* = 2.1 Hz, 1H), 8.22 – 8.13 (m, 2H), 8.07 (t, *J* = 8.6 Hz, 3H), 7.86 (d, *J* = 6.0 Hz, 1H), 7.31 (t, *J* = 7.8 Hz, 3H), 7.10 – 7.03 (m, 3H), 6.84 – 6.77 (m, 3H), 6.72 (d, *J* = 1.9 Hz, 1H), 6.61 (s, 1H), 6.53 (s, 2H), 5.55 – 5.46 (m, 1H), 5.27 (s, 1H), 4.03 – 3.96 (m, 3H), 3.89 (s, 2H), 3.70 (d, *J* = 4.0 Hz, 7H), 3.61 – 3.52 (m, 19H), 3.48 (d, *J* = 6.6 Hz, 5H), 2.57 (d, *J* = 4.5 Hz, 6H), 1.98 – 1.84 (m, 4H), 1.58 (dd, *J* = 16.8, 10.2 Hz, 5H), 0.80 (t, *J* = 7.2 Hz, 3H). ESI [M+H]^+^: 1248.3

**<u>Pan KRAS inhibitor</u>** (Yamano, M. M.; Li, Y.; Navaratne, P.; Medina, J.; Chen, N.; Pettus, L.; Rahimoff, R.; Li, X.; Stellwagen, J.; Manoni, F.; Li, K.; Lanman, B.A.; Wurz, R.P.; Zhao, W.; Rui, H.; Eshon, J. Heterocyclic Compounds and Methods of Use. WO 2023/018812 A1, 2023).

**2-(8-Ethyl-7-fluoro-3-(methoxymethoxy)naphthalen-1-yl)-4,4,5,5-tetramethyl-1,3,2-dioxaborolane (Intermediate A)**

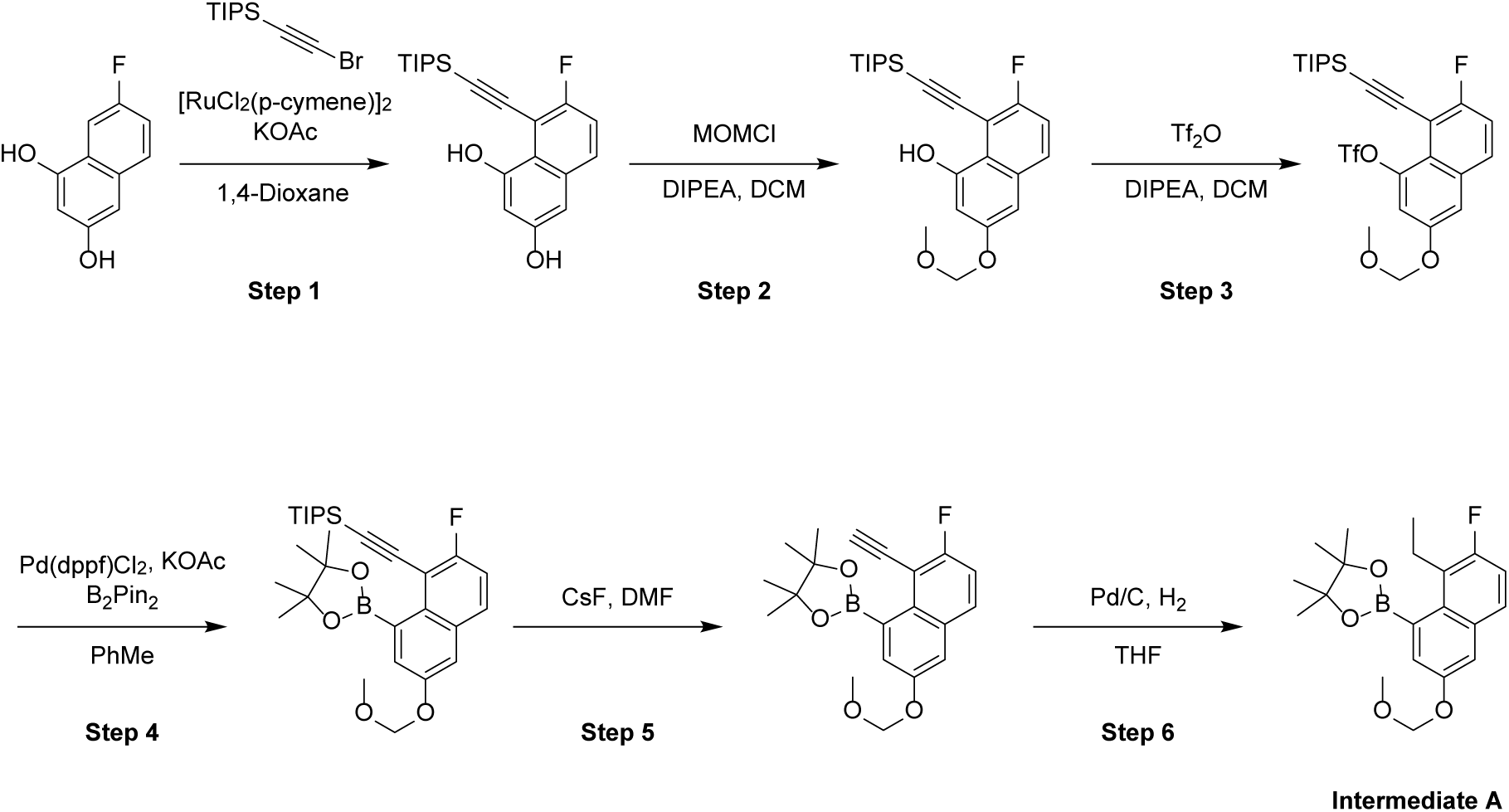

**Step-1: Synthesis of 7-Fluoro-8-((triisopropylsilyl)ethynyl) naphthalene-1,3-diol.**

To a mixture of 7-fluoronaphthalene-1,3-diol (3.30 g, 18.52 mmol) and 2-bromoethynyl(triisopropyl)silane (5.81 g, 22.23 mmol) in dioxane (60 mL) was added KOAc (3.64 g, 37.05 mmol) and dichloro(*p*-cymene) ruthenium(II) dimer (1.13 g, 1.85 mmol, 0.1 equiv.) in one portion at 15 °C under nitrogen. The mixture was stirred at 110 °C for 2 h. The mixture was cooled to 15 °C and poured into ice-water (60 mL). The mixture was extracted with ethyl acetate (3 x 50 mL). The combined organic phases were dried over anhydrous Na2SO4, filtered and concentrated in vacuo. The crude residue was purified by column chromatography on silica gel, eluting with petroleum ether/ethyl acetate 8/1 to 1/1 to give 7-fluoro-8-((triisopropylsilyl)ethynyl) naphthalene-1,3-diol (4.30 g, 11.27 mmol, 65% yield) as a yellow solid.

ESI [M+H]^+^: 359.2

**Step-2: Synthesis of 7-Fluoro-3-(methoxymethoxy)-8-((triisopropylsilyl)ethynyl)naphthalen-1-ol.**

To a mixture of 7-fluoro-8-((triisopropylsilyl)ethynyl)naphthalene-1,3-diol (4.30 g, 11.99 mmol) in DCM (60 mL) was added DIPEA (4.65 g, 35.98 mmol, 6.27 mL). Then MOM-Cl (1.16 g, 14.39 mmol, 1.09 mL) was added in portions at 0 °C under nitrogen. The mixture was stirred at 15 °C for 12 h. The mixture was poured into ice-water (60 mL) and stirred for 20 min. The mixture was extracted with ethyl acetate (3 x 60 mL). The combined organic phase was washed with brine (2 x 60 mL), dried over anhydrous Na2SO4, filtered and concentrated. The residue was purified by column chromatography on silica gel, eluting with petroleum ether/ethyl acetate 4/1 to 1/1 to give 7-fluoro-3-(methoxymethoxy)-8-(2-triisopropylsilylethynyl)naphthalen-1-ol (2.00 g, 4.97 mmol, 41% yield) as a brown solid.

**Step-3: Synthesis of 7-Fluoro-3-(methoxymethoxy)-8-((triisopropylsilyl)ethynyl)naphthalen-1-yl trifluoromethanesulfonate.**

To a mixture of 7-fluoro-3-(methoxymethoxy)-8-(2-triisopropylsilylethynyl)naphthalen-1-ol (2.50 g, 6.21 mmol) in DCM (30 mL) was added DIPEA (2.41 g, 18.63 mmol, 3.25 mL) in one portion at 15 °C under nitrogen. Then to the mixture was added Tf2O (2.63 g, 9.32 mmol, 1.54 mL) in portions at -40 °C under nitrogen. The mixture was stirred at -40 °C for 1 h. The mixture was poured into ice-water (30 mL) and the aqueous phase was extracted with dichloromethane (3 x 30 mL). The combined organic phase was dried with anhydrous Na2SO4, filtered and concentrated in vacuo. The residue was purified by column chromatography on silica gel, eluting with petroleum ether/ethyl acetate 40/1 to 8/1 to give 7-fluoro-3-(methoxymethoxy)-8-((triisopropylsilyl)ethynyl)naphthalen-1-yl trifluoromethanesulfonate (2.50 g, 4.68 mmol, 75% yield) as ayellow oil.

**Step-4: Synthesis of ((2-Fluoro-6-(methoxymethoxy)-8-(4,4,5,5-tetramethyl-1,3,2-dioxaborolan-2-yl)naphthalen-1-yl)ethynyl)triisopropylsilane.**

To a mixture of 7-fluoro-3-(methoxymethoxy)-8-((triisopropylsilyl)ethynyl)naphthalen-1-yl trifluoromethanesulfonate (2.50 g, 4.68 mmol) in toluene (25 mL) was added KOAc (1.38 g, 14.03 mmol), bis(pinacolato)diboron (2.37 g, 9.35 mmol) and Pd(dppf)Cl2 (0.34 g, 0.47 mmol) in one portion at 15 °C under nitrogen. The mixture was stirred at 130 °C for 3 h then cooled to 15 °C and concentrated under reduced pressure at 45 °C. The residue was poured into ice-water (40 mL). The mixture was extracted with ethyl acetate (3 x 40 mL) and the combined organic phase was dried over anhydrous Na2SO4, filtered and concentrated in vacuo. The residue was purified by column chromatography on silica gel, eluting with petroleum ether/ethyl acetate 40/1 to 10/1 to give ((2-fluoro-6-(methoxymethoxy)-8-(4,4,5,5-tetramethyl-1,3,2-dioxaborolan-2-yl)naphthalen-1-yl)ethynyl)triisopropylsilane (1.20 g, 2.34 mmol, 50% yield) as a yellow solid. ESI [M+H]^+^: 513.2

**Step-5: Synthesis of 2-(8-Ethynyl-7-fluoro-3-(methoxymethoxy)naphthalen-1-yl)-4,4,5,5-tetramethyl-1,3,2-dioxaborolane.**

To a mixture of ((2-fluoro-6-(methoxymethoxy)-8-(4,4,5,5-tetramethyl-1,3,2-dioxaborolan-2-yl)naphthalen-1-yl)ethynyl)triisopropylsilane (1.10 g, 2.15 mmol) in DMF (20 mL) was added CsF (1.96 g, 12.88 mmol) in one portion at 15 °C under nitrogen. The mixture was stirred at 50 °C for 2 h then the mixture was poured into water (20 mL). The mixture was extracted with ethyl acetate (3 x 20 mL). The combined organic phase was washed with brine (2 x 30 mL), dried over anhydrous Na2SO4, filtered and concentrated in vacuo to give the crude 2-(8-ethynyl-7-fluoro-3-(methoxymethoxy)naphthalen-1-yl)-4,4,5,5-tetramethyl-1,3,2-dioxaborolane (1.20 g, crude) as a brown oil.

**Step-6: Synthesis of 2-(8-Ethyl-7-fluoro-3-(methoxymethoxy)naphthalen-1-yl)-4,4,5,5-tetramethyl-1,3,2-dioxaborolane (Intermediate A).**

To a solution of 2-(8-ethynyl-7-fluoro-3-(methoxymethoxy)naphthalen-1-yl)-4,4,5,5-tetramethyl-1,3,2-dioxaborolane (1.20 g, 3.37 mmol) in THF (10 mL) was added Pd/C (10%, 20 mg) under argon. The suspension was degassed under vacuum and purged with hydrogen several times. The mixture was stirred under hydrogen (15 psi) at 15 °C for 1 h. The reaction mixture was filtered, and the filter cake was washed with THF (3 x 10 mL). The combined filtrate was concentrated under reduced pressure. The crude product was purified by column chromatography on silica gel, eluting with petroleum ether/ethyl acetate 25/1 to 10/1 to give 2-(8-ethyl-7-fluoro-3-(methoxymethoxy)naphthalen-1-yl)-4,4,5,5-tetramethyl-1,3,2-dioxaborolane (**Intermediate A**, 0.62 g, 1.69 mmol, 75% yield over two steps) as a yellow solid. ESI [M+H]^+^: 361.1

**(*R*)-1-(8-Fluoro-2-(((2*R*,7a*S*)-2-fluorotetrahydro-1*H*-pyrrolizin-7a(5*H*)-yl)methoxy)-7-(tributylstannyl)pyrido[4,3-*d*]pyrimidin-4-yl)-3-methylpiperidin-3-ol (Intermediate B).**

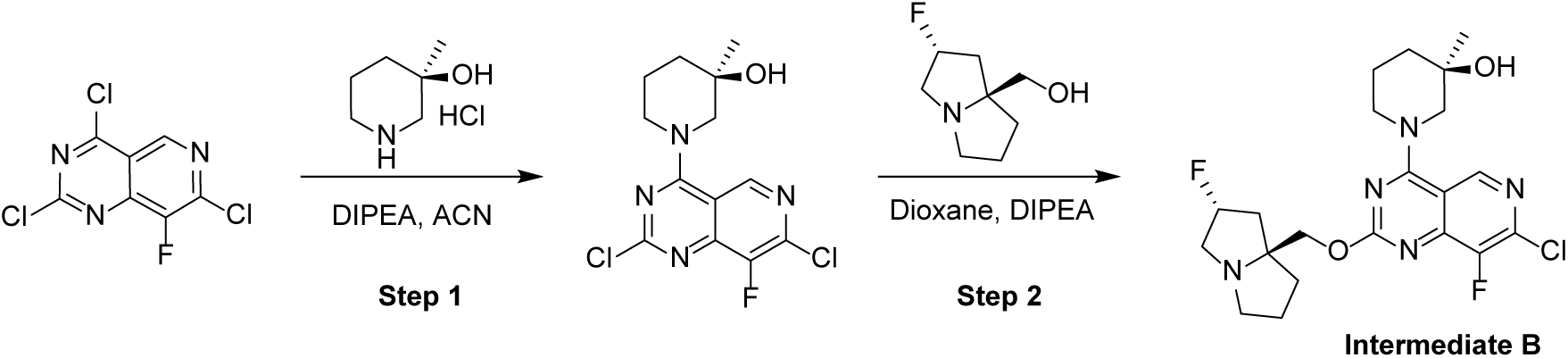

**Step-1: Synthesis of (*R*)-1-(2,7-Dichloro-8-fluoropyrido[4,3-*d*]pyrimidin-4-yl)-3-methylpiperidin-3-ol.**

To a mixture of 2,4,7-trichloro-8-fluoropyrido[4,3-*d*]pyrimidine (25 g, 99 mmol) in acetonitrile (500 mL) was added DIPEA (86 mL, 495 mmol) and (*R*)-3-methylpiperidin-3-ol hydrochloride (15 g, 99 mmol). The mixture was stirred at 0 °C for 0.5 h under nitrogen. The reaction mixture was diluted with H2O (1 L) and extracted with EtOAc (3 x 0.9 L). The combined organic layers were dried over Na2SO4, filtered and the filtrate was concentrated under reduced pressure. The residue was purified by chromatography on silica gel, eluting with a gradient of 2-100% ethyl acetate/petroleum ether to give (*R*)-1-(2,7-dichloro-8-fluoropyrido[4,3-*d*]pyrimidin-4-yl)-3-methylpiperidin-3-ol (28 g, 85 mmol, 85% yield) as a yellow solid.

**Step-2: Synthesis of (*R*)-1-(7-Chloro-8-fluoro-2-(((2*R*,7a*S*)-2-fluorohexahydro-1*H*-pyrrolizin-7a-yl)methoxy)pyrido[4,3-*d*]pyrimidin-4-yl)-3-methylpiperidin-3-ol.**

To a solution of (*R*)-1-(2,7-dichloro-8-fluoropyrido[4,3-*d*]pyrimidin-4-yl)-3-methylpiperidin-3-ol (20 g, 60 mmol) in 1,4-dioxane (300 mL) was added DIPEA (20 g, 151 mmol) and ((2*R*,7a*S*)-2-fluorotetrahydro-1*H*-pyrrolizin-7a(5*H*)-yl)methanol (14 g, 85 mmol). The resulting mixture was stirred at 100 °C for 12 h under nitrogen. The reaction mixture was diluted with H2O (0.7 L) and extracted with EtOAc (3 x 0.5 L). The combined organic layers were dried over Na2SO4, filtered and concentrated. The residue was purified by column chromatography on silica gel, eluting with a gradient of 10-100% ethyl acetate/petroleum ether to give (*R*)-1-(7-chloro-8-fluoro-2-(((2*R*,7a*S*)-2-fluorohexahydro-1*H*-pyrrolizin-7a-yl)methoxy)pyrido[4,3-*d*]pyrimidin-4-yl)-3-methylpiperidin-3-ol (**Intermediate B**). Batch 1: 10.3 g with 98.5% purity by LCMS; Batch 2: 3.3 g crude with 70% purity. ESI [M+H]^+^: 454.2, 456.2

**(*R*)-1-(7-(8-ethyl-7-fluoro-3-hydroxynaphthalen-1-yl)-8-fluoro-2-(((2*R*,7a*S*)-2-fluorotetrahydro-1*H*-pyrrolizin-7a(5*H*)-yl)methoxy)pyrido[4,3-*d*]pyrimidin-4-yl)-3-methylpiperidin-3-ol (Pan KRAS inhibitor).**

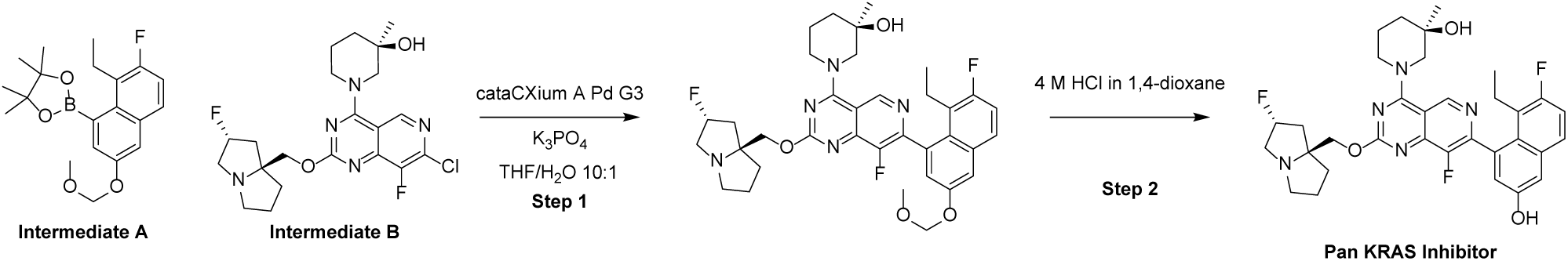

**Step-1: Synthesis of (*R*)-1-(7-(8-ethyl-7-fluoro-3-(methoxymethoxy)naphthalen-1-yl)-8-fluoro-2-(((2*R*,7a*S*)-2-fluorotetrahydro-1*H*-pyrrolizin-7a(5*H*)-yl)methoxy)pyrido[4,3-*d*]pyrimidin-4-yl)-3-methylpiperidin-3-ol.**

To a solution of 2-(8-ethyl-7-fluoro-3-(methoxymethoxy)naphthalen-1-yl)-4,4,5,5-tetramethyl-1,3,2-dioxaborolane (**Intermediate A**, 4.29 g, 11.90 mmol) and (*R*)-1-(7-chloro-8-fluoro-2-(((2*R*,7a*S*)-2-fluorotetrahydro-1*H*-pyrrolizin-7a(5*H*)-yl)methoxy)pyrido[4,3-*d*]pyrimidin-4-yl)-3-methylpiperidin-3-ol (**Intermediate B**, 3.6 g, 7.93 mmol) in tetrahydrofuran (72 mL) and water (7.2 mL) were added potassium phosphate (5.05 g, 23.79 mmol) and cataCXium A Pd G3 (0.58 g, 0.79 mmol, Sigma Aldrich). The reaction mixture was sparged with argon, sealed, and stirred at 70 °C for 16 h. The reaction mixture was concentrated and partitioned between water and ethyl acetate; the organic layer was concentrated. The crude product was purified by column chromatography on silica gel, eluting with 0-100% (3:1 ethyl acetate/ethanol + 2% TEA) in heptane, to provide (*R*)-1-(7-(8-ethyl-7-fluoro-3-(methoxymethoxy)naphthalen-1-yl)-8-fluoro-2-(((2*R*,7a*S*)-2-fluorotetrahydro-1*H*-pyrrolizin-7a(5*H*)-yl)methoxy)pyrido[4,3-*d*]pyrimidin-4-yl)-3-methylpiperidin-3-ol (5.15 g, 7.90 mmol, quantitative yield) as a yellow powder. ESI [M+H]^+^: 652.3

**Step-2: Synthesis of (*R*)-1-(7-(8-Ethyl-7-fluoro-3-hydroxynaphthalen-1-yl)-8-fluoro-2-(((2*R*,7a*S*)-2-fluorotetrahydro-1*H*-pyrrolizin-7a(5*H*)-yl)methoxy)pyrido[4,3-*d*]pyrimidin-4-yl)-3-methylpiperidin-3-ol.**

To a solution of (*R*)-1-(7-(8-ethyl-7-fluoro-3-(methoxymethoxy)naphthalen-1-yl)-8-fluoro-2-(((2*R*,7a*S*)-2-fluorotetrahydro-1*H*-pyrrolizin-7a(5*H*)-yl)methoxy)pyrido[4,3-*d*]pyrimidin-4-yl)-3-methylpiperidin-3-ol (5.00 g, 7.67 mmol) in acetonitrile (38.4 mL) was added hydrogen chloride solution (4.0 M in 1,4-dioxane, 19.18 mL, 77 mmol). The reaction was stirred at ambient temperature for 30 min then MeOH (∼5 mL) was added to quench any reactive intermediates, and the reaction mixture was concentrated. The crude product was purified by column chromatography on silica gel, eluting with 0-100% (3:1 ethyl acetate/ethanol + 2% TEA) in heptane, to provide (*R*)-1-(7-(8-ethyl-7-fluoro-3-hydroxynaphthalen-1-yl)-8-fluoro-2-(((2*R*,7a*S*)-2-fluorotetrahydro-1*H*-pyrrolizin-7a(5*H*)-yl)methoxy)pyrido[4,3-*d*]pyrimidin-4-yl)-3-methylpiperidin-3-ol (3.36 g, 5.53 mmol, 72% yield) as a yellow powder. ^1^H NMR (400 MHz*, METHANOL-d4*): δ 9.22 (s, 1 H), 7.69 (dd, *J*=8.9, 5.7 Hz, 1 H), 7.23 - 7.33 (m, 2 H), 7.08 (s, 1 H), 5.18 - 5.44 (m, 1 H), 4.52 (br d, *J*=13.0 Hz, 1 H), 4.25 - 4.36 (m, 3 H), 3.59 - 3.70 (m, 1 H), 3.48 - 3.54 (m, 1 H), 3.15 - 3.27 (m, 3 H), 2.99 - 3.06 (m, 1 H), 2.49 (br dd, *J*=14.0, 6.9 Hz, 1 H), 2.10 - 2.30 (m, 5 H), 1.76 - 2.05 (m, 8 H), 1.30 (d, *J*=9.4 Hz, 3 H), 0.83 (q, *J*=7.7 Hz, 3 H). ^19^F NMR (376 MHz, *METHANOL-d4*) δ ppm 121.2 (s, 1 F), 139.1 (s, 1 F), 173.7 (s, 1 F). ESI [M+H]^+^: 608.2

### KRAS PROTAC

**(2S,4R)-1-((S)-2-(4-((((2S,7aR)-7a-(((7-(8-ethyl-7-fluoro-3-hydroxynaphthalen-1-yl)-8-fluoro-4-((R)-3-hydroxy-3-methylpiperidin-1-yl)pyrido[4,3-d]pyrimidin-2-yl)oxy)methyl)hexahydro-1H-pyrrolizin-2-yl)oxy)methyl)-1H-1,2,3-triazol-1-yl)-3,3-dimethylbutanoyl)-4-hydroxy-N-((S)-1-(4-(4-methylthiazol-5-yl)phenyl)ethyl)pyrrolidine-2-carboxamide**

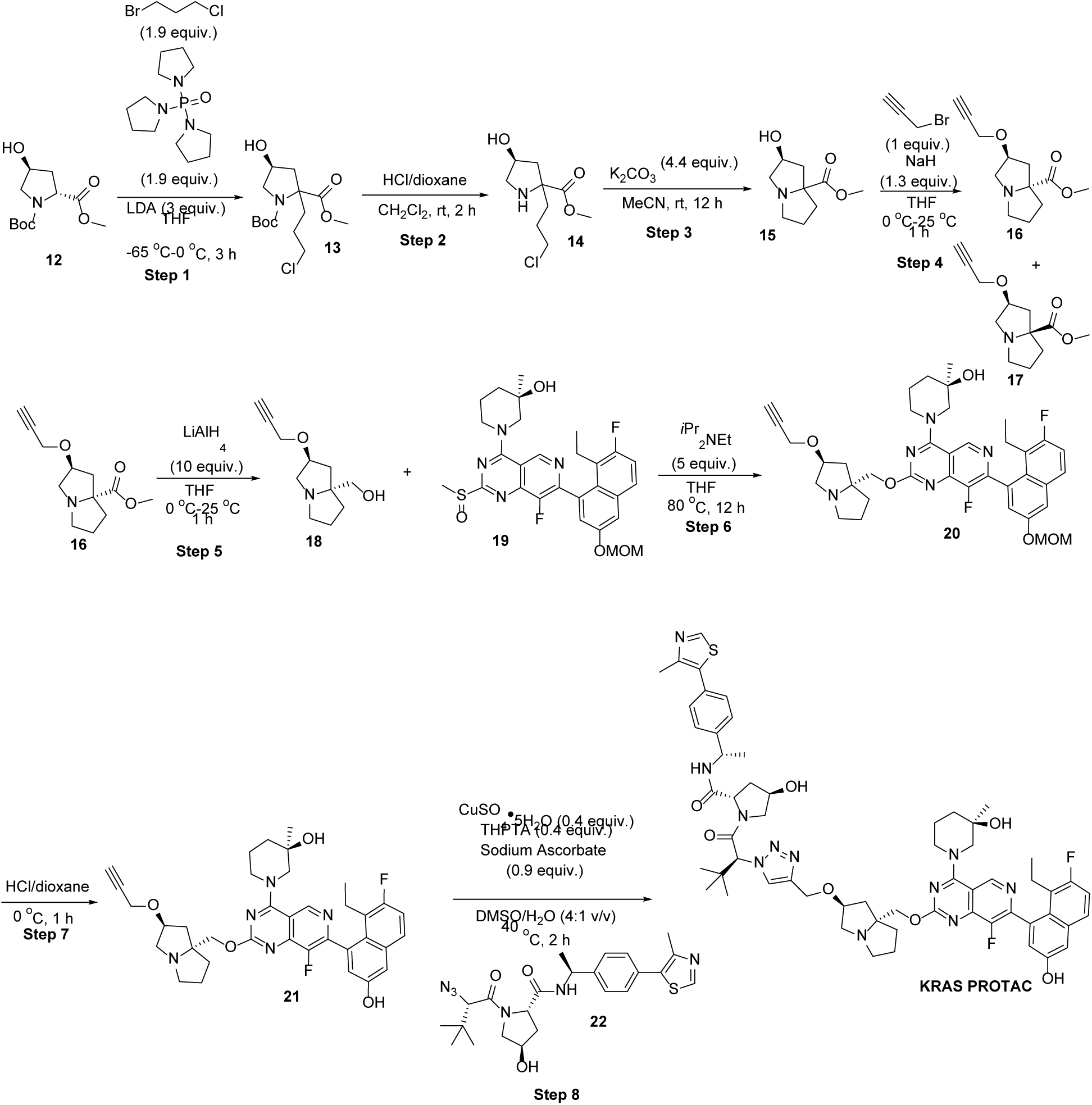

**Step 1: 1-(tert-butyl) 2-methyl (4S)-2-(3-chloropropyl)-4-hydroxypyrrolidine-1,2-dicarboxylate**

To a solution of LDA (673 mL, 1345 mmol,1M in THF) in tetrahydrofuran (611 mL) was added tri(pyrrolidin-1-yl)phosphine oxide (219 g, 852 mmol) and 1-(tert-butyl) 2-methyl (2R,4S)-4-hydroxypyrrolidine-1,2-dicarboxylate (110 g, 448 mmol) in tetrahydrofuran (160 mL) at -65 °C. The mixture was stirred at -65°C for 10 min. Then it was stirred at 0 °C for 1 h. Compound1-bromo-3-chloropropane (134 g, 852 mmol) was added the above mixture at -65 °C. The mixture was stirred at 25 °C for 2 h. TLC (petroleum ether: ethyl acetate = 1:1, product: Rf = 0.35) showed was consumed completely and one main new spot was formed. The reaction mixture was quenched by addition NH4Cl (300 mL), and extracted with ethyl acetate (2 × 300 mL). The combined organic layers were washed with brine (300 mL), dried over Na2SO4, filtered and concentrated under reduced pressure to give a residue. Combined with another batch (90 g scale). The residue was purified by column chromatography (SiO2, petroleum ether/ethyl acetate=100/1 to 1/1, TLC showed (petroleum ether: ethyl acetate = 1/1, product: Rf = 0.35) to obtain 1-(tert-butyl) 2-methyl (4S)-2-(3-chloropropyl)-4-hydroxypyrrolidine-1,2-dicarboxylate (208 g, 647 mmol, 79% yield) as a yellow oil.

**Step-2: (4S)-2-(3-chloropropyl)-4-hydroxypyrrolidine-2-carboxylate**

To a solution of 1-(tert-butyl) 2-methyl (4S)-2-(3-chloropropyl)-4-hydroxypyrrolidine-1,2-dicarboxylate (50 g, 155 mmol) in dichloromethane (360 mL) was added HCl-Dioxane (252 mL, 1010 mmol, 4M). Then the mixture was stirred at 25 °C for 2 h. TLC (dichloromethane: methanol = 10:1) was consumed completely and one main new spot was formed. The reaction mixture was concentrated under reduced pressure to give compound methyl (4S)-2-(3-chloropropyl)-4-hydroxypyrrolidine-2-carboxylate (40 g, 180 mmol, HCl salt, 99% yield).

**Step-3: (2S)-2-hydroxytetrahydro-1H-pyrrolizine-7a(5H)-carboxylate**

To a solution of methyl (4S)-2-(3-chloropropyl)-4-hydroxypyrrolidine-2-carboxylate (40 g, 180 mmol) in acetonitrile (280 mL) was added K2CO3 (100 g, 722 mmol). Then the mixture was stirred at 25 °C for 12 h. TLC (dichloromethane: methanol = 5:1, product: Rf = 0.2) showed was consumed completely and one main new spot was formed. The reaction mixture was filtered, the filtrate was concentrated in vacuum to afford compound methyl (2S)-2-hydroxytetrahydro-1H-pyrrolizine-7a(5H)-carboxylate (22 g, 119 mmol, 66% yield) as a brown oil.

**Step-4: methyl (2S,7aR)-2-(prop-2-yn-1-yloxy)tetrahydro-1H-pyrrolizine-7a(5H)-carboxylate**

To a solution of methyl (2S)-2-hydroxytetrahydro-1H-pyrrolizine-7a(5H)-carboxylate (22 g, 119 mmol) in tetrahydrofuran (130 mL) was added NaH (6.18 g, 154 mmol, 60% purity) in portions at 0 °C and stirred for 0.5 h. The mixture was cooled to 0 °C, then 3-bromoprop-1-yne (16.96 g, 143 mmol) was added to the mixture at 0 °C, and then the mixture was warmed to 25 °C and stirred for 1 h. TLC (dichloromethane : methanol = 8:1) indicated starting material was consumed completely and two new spots formed (Rf =0.6, 0.4). Combined with other 3 batches (20 g scale). The combined reaction mixture was filtered and concentrated under reduced pressure to give a residue. The residue was purified by column chromatography (SiO2, Petroleum ether/Ethyl acetate =30 /1 to 0 /1) to obtain Spot 1: (Rf =0.6) methyl (2S,7aR)-2-(prop-2-yn-1-yloxy)tetrahydro-1H-pyrrolizine-7a(5H)-carboxylate **S5** (10 g, 44 mmol, 13% yield) as a yellow oil, and Spot 2: (Rf =0.4) methyl (2S,7aS)-2-(prop-2-yn-1-yloxy)tetrahydro-1H-pyrrolizine-7a(5H)-carboxylate **S6** (3.3 g, 14 mmol, 4% yield) was obtained as a yellow oil.

**Step-5: ((2S,7aR)-2-(prop-2-yn-1-yloxy)tetrahydro-1H-pyrrolizin-7a(5H)-yl)methanol**

To a mixture of LAH (3.58 ml, 2.5 M) in THF (10 mL) was added methyl (2S,7aR)-2-(prop-2-yn-1-yloxy)tetrahydro-1H-pyrrolizine-7a(5H)-carboxylate (2 g, 8.96 mmol) in THF (7 mL) dropwise at 0 °C. The mixture was warmed to 25 °C and stirred for 1 h. TLC (dichloromethane: methanol = 1/1, Rf = 0.1) showed the reaction was completed. The mixture was quenched with water (2 × 5 mL) and 15% NaOH solution (2 × 5 mL). Total five batches were carried out in parallel and combined for the work up. The mixture was filtered and the filtrate was concentrated under vacuum.

((2S,7aR)-2-(prop-2-yn-1-yloxy)tetrahydro-1H-pyrrolizin-7a(5H)-yl)methanol (8.5g, 4.51mmol, 91 % yield) was obtained as a yellow oil. ^1^H NMR (400 MHz, CHLOROFORM-*d*) δ 4.33 - 4.29 (m, 1H), 4.13 - 4.10 (m, 2H), 3.47 (s, 2H), 3.25 - 3.20 (m, 3H), 3.05 – 2.95 (m, 1H), 2.91 - 2.75 (m, 2H), 2.15 - 2.10 (m, 1H), 1.90 -1.80 (m, 1H), 1.80 - 1.60 (m, 4H).

**Step 6: (R)-1-(7-(8-ethyl-7-fluoro-3-(methoxymethoxy)naphthalen-1-yl)-8-fluoro-2-(((2S,7aR)-2-(prop-2-yn-1-yloxy)tetrahydro-1H-pyrrolizin-7a(5H)-yl)methoxy)pyrido[4,3-d]pyrimidin-4-yl)-3-methylpiperidin-3-ol**

To a solution of ((2S,7aR)-2-(prop-2-yn-1-yloxy)tetrahydro-1H-pyrrolizin-7a(5H)-yl)methanol (2 g, 10.24 mmol) in tetrahydrofuran (20 mL) was added (3R)-1-(7-(8-ethyl-7-fluoro-3-(methoxymethoxy)naphthalen-1-yl)-8-fluoro-2-(methylsulfinyl)pyrido[4,3-d]pyrimidin-4-yl)-3-methylpiperidin-3-ol (5.70 g, 10.24 mmol) (synthesized using procedure outlined in WO20230188A1) and *i*Pr2NEt (8.94 mL, 51.2 mmol). Then the mixture was stirred at 80°C for 8 h under N2 atmosphere. TLC (DCM: MeOH = 10:1) indicated starting material was consumed completely and main one spot formed (Rf = 0.5). The reaction mixture was diluted with water (30 mL) and extracted with EtOAc (3 × 20 mL). The combined organic layers were washed with brine (3 × 15 mL), dried over Na2SO4, filtered and concentrated under reduced pressure to give a residue. The residue was purified by prep-HPLC (column: Phenomenex Luna C18 250mm × 100mm × 10um;mobile phase: [A: H2O(10mM NH4HCO3);B: ACN];B%: 40.00%-70.00%,20.00min). Compound (R)-1-(7-(8-ethyl-7-fluoro-3-(methoxymethoxy)naphthalen-1-yl)-8-fluoro-2-(((2S,7aR)-2-(prop-2-yn-1-yloxy)tetrahydro-1H-pyrrolizin-7a(5H)-yl)methoxy)pyrido[4,3-d]pyrimidin-4-yl)-3-methylpiperidin-3-ol (1.68 g, 2.443 mmol, 24 % yield) was obtained as a yellow solid.

ESI [M+H]^+^: 688.3.

**Step 7: (R)-1-(7-(8-ethyl-7-fluoro-3-hydroxynaphthalen-1-yl)-8-fluoro-2-(((2S,7aR)-2-(prop-2-yn-1-yloxy)tetrahydro-1H-pyrrolizin-7a(5H)-yl)methoxy)pyrido[4,3-d]pyrimidin-4-yl)-3-methylpiperidin-3-ol**

To a solution of (R)-1-(7-(8-ethyl-7-fluoro-3-(methoxymethoxy)naphthalen-1-yl)-8-fluoro-2-(((2R,7aR)-2-(prop-2-yn-1-yloxy)tetrahydro-1H-pyrrolizin-7a(5H)-yl)methoxy)pyrido[4,3-d]pyrimidin-4-yl)-3-methylpiperidin-3-ol (0.2 g, 0.291 mmol) in dichloromethane (6 mL) was added HCl-Dioxane (0.4 mL, 4 M) in sequence. Then the mixture was stirred at 0 °C for 1 h. LCMS showed one main peak with desired m/z was detected. Total 5 batches were carried out in parallel and combined for the work up. The mixture was concentrated to get a residue. The residue was purified by prep-HPLC. (R)-1-(7-(8-ethyl-7-fluoro-3-hydroxynaphthalen-1-yl)-8-fluoro-2-(((2R,7aR)-2-(prop-2-yn-1-yloxy)tetrahydro-1H-pyrrolizin-7a(5H)-yl)methoxy)pyrido[4,3-d]pyrimidin-4-yl)-3-methylpiperidin-3-ol (47.8 mg, 0.074 mmol, 12.7%) was obtained as a white solid. ^1^H NMR (400 MHz, CHLOROFORM- *d*) δ 9.20 - 9.00 (m, 1H), 7.56 - 7.51 (m, 1H), 7.21 - 7.14 (m, 2H), 7.15 – 6.75 (m, 1H), 4.50 - 4.30 (m, 3H), 4.30 - 4.20 (m, 2H), 4.15 - 4.10 (m, 2H), 3.68 - 3.10 (m, 5H), 2.92 - 2.82 (m, 1H), 2.78 - 2.61 (m, 1H), 2.52 - 2.40 (m, 1H), 2.39 - 2.30 (m, 2H), 2.28 - 2.08 (m, 3H), 2.00 – 1.40 (m, 8H), 1.28 - 1.23 (m, 3H), 0.84 - 0.75 (m, 3H). ESI [M+H]^+^: 644.5.

**Step 8: (2S,4R)-1-((S)-2-(4-((((2S,7aR)-7a-(((7-(8-ethyl-7-fluoro-3-hydroxynaphthalen-1-yl)-8- fluoro-4-((R)-3-hydroxy-3-methylpiperidin-1-yl)pyrido[4,3-d]pyrimidin-2-yl)oxy)methyl)hexahydro-1H-pyrrolizin-2-yl)oxy)methyl)-1H-1,2,3-triazol-1-yl)-3,3-dimethylbutanoyl)-4-hydroxy-N-((S)-1-(4-(4-methylthiazol-5-yl)phenyl)ethyl)pyrrolidine-2-carboxamide (KRAS PROTAC)**

To a solution of (R)-1-(7-(8-ethyl-7-fluoro-3-hydroxynaphthalen-1-yl)-8-fluoro-2-(((2S,7aR)-2-(prop-2-yn-1-yloxy)tetrahydro-1H-pyrrolizin-7a(5H)-yl)methoxy)pyrido[4,3-d]pyrimidin-4-yl)-3-methylpiperidin-3-ol (539 mg, 0.837 mmol)in DMSO (3.12 mL) and water (0.32 mL) was added (2S,4R)-1-((S)-2-azido-3,3-dimethylbutanoyl)-4-hydroxy-N-((S)-1-(4-(4-methylthiazol-5-yl)phenyl)ethyl)pyrrolidine-2-carboxamide (402 mg, 0.854 mmol) followed by a pre-mixed solution of copper (II) sulfate pentahydrate (84 mg, 0.335 mmol) and tris(3-hydroxypropyltriazolylmethyl)amine (145 mg, 0.335 mmol). The reaction mixture was stirred at 40 °Cfor 1.5 h. The reaction mixture was cooled to rt. The reaction mixture was diluted with saturated NaCl solution and extracted with 2-Me-THF (2 x 50 mL). The organic extract was washed with saturatedd NaCl solution (20 mL) and dried over MgSO4. The solution was filtered and concentrated *in vacuo* to give the crude material which was absorbed onto a plug of silica gel and purified by chromatography through a Redi-Sep pre-packed silica gel column (80 g) eluting with a gradient of0 % to 80 % MeOH in CH2Cl2, to provide (2S,4R)-1-((S)-2-(4-((((2S,7aR)-7a-(((7-(8-ethyl-7-fluoro-3-hydroxynaphthalen-1-yl)-8-fluoro-4-((R)-3-hydroxy-3-methylpiperidin-1-yl)pyrido[4,3-d]pyrimidin-2-yl)oxy)methyl)hexahydro-1H-pyrrolizin-2-yl)oxy)methyl)-1H-1,2,3-triazol-1-yl)-3,3-dimethylbutanoyl)-4-hydroxy-N-((S)-1-(4-(4-methylthiazol-5-yl)phenyl)ethyl)pyrrolidine-2-carboxamide (290 mg, 0.260 mmol, 31 % yield) as an off-white solid after filtration over 0.45 um filter. ^1^H NMR (500 MHz, DMSO-d^6^) δ 10.0-9.8 (m, 1H), 9.30-9.10 (m, 1H), 9.00-8.90 (m, 1H), 8.60-8.40 (m, 1H), 8.30-8.20 (m, 1H), 7.90-7.70 (m, 1H), 7.50-7.30 (m, 6H), 7.10-7.00 (m, 1H), 5.60-5.40 (m, 1H), 5.20-5.10 (m, 1H), 5.00-4.80 (m, 1H), 4.78-4.70 (m, 1H), 4.60-4.10 (m, 6H), 4.08-4.00 (m, 3H), 3.70-3.50 (m, 3H), 3.50-3.40 (m, 1H), 3.10-2.90 (m, 2H), 2.88-2.70 (m, 2H), 2.50-2.40 (m, 3H), 2.40-2.30 (m, 1H), 2.20-2.00 (m, 4H), 1.98-1.80 (m, 2H), 1.78-1.60 (m, 7H), 1.50-1.40 (m, 3H), 1.20-1.10 (m, 3H), 1.00-0.90 (m, 9H), 0.80-0.70 (m, 3H). ESI [M+H]^+^: 1114.5.

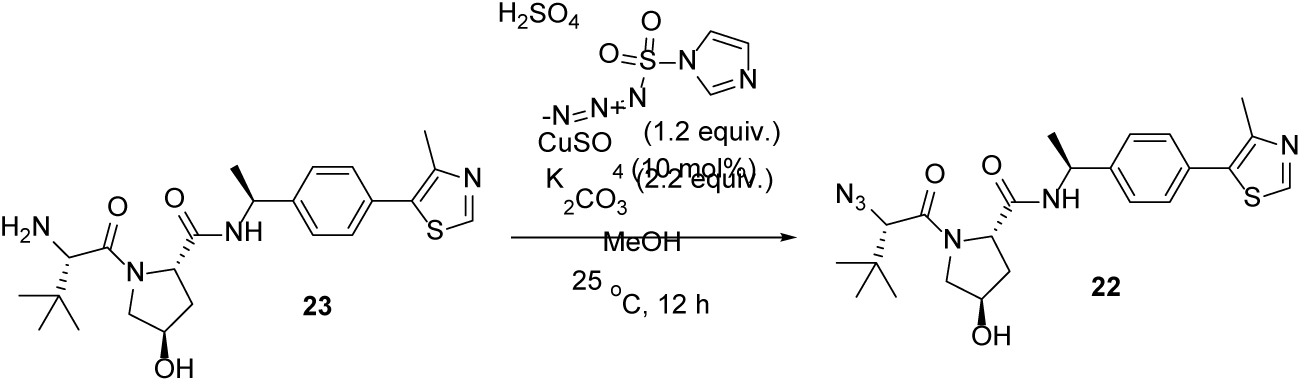

To a suspension of (2S,4R)-1-((S)-2-amino-3,3-dimethylbutanoyl)-4-hydroxy-N-((S)-1-(4-(4-methylthiazol-5-yl)phenyl)ethyl)pyrrolidine-2-carboxamide (1.5 g, 3.37 mmol) and potassium carbonate (1.026 g, 7.42 mmol) in methanol (16.87 mL) followed by copper(II) sulfate anhydrous (0.054 g, 0.337 mmol) and 1-(azidosulfonyl)-1h-imidazol-3-ium hydrogensulfate (1.098 g, 4.05 mmol) The reaction mixturewas stirred at 25 °Cfor 2 h (minimal conversion to the desired product m/z at this point). Additional K2CO3 (1.026 g, 7.42 mmol, 2 equiv.) was added and the reaction mixture was stirred at rt for another 12 h. The reaction mixture was diluted with saturated NaHCO3 solution (10 mL) and extracted with CH2Cl2 (2 x 20 mL). The organic extract was washed with saturated NaCl solution (10 mL) and dried over MgSO4. The solution was filtered and concentrated *in vacuo* to give the crude material which was absorbed onto a plug of silica gel and purified by chromatography through a Redi-Sep pre-packed silica gel column (80 g), eluting with a gradient of 0 % to 10 % MeOH in CH2Cl2 to provide (2S,4R)-1-((S)-2-azido-3,3-dimethylbutanoyl)-4-hydroxy-N-((S)-1-(4-(4-methylthiazol-5-yl)phenyl)ethyl)pyrrolidine-2-carboxamide (0.678 g, 1.441 mmol, 43 % yield) as pale yellow solid. ^1^H NMR (400 MHz, CHLOROFORM-*d*) δ ppm 8.74 - 8.60 (m, 1H), 7.64 - 7.56 (m, 1H), 7.46 -7.36 (m, 4H), 5.13 - 5.01 (m, 1H), 4.94 - 4.74 (m, 1H), 4.71 - 4.53 (m, 1H), 3.76 - 3.56 (m, 3H), 2.84 - 2.68 (m, 1H), 2.59 - 2.43 (m, 4H), 2.16 – 1.98 (m, 1H), 1.57 - 1.44 (m, 3H), 1.14 - 1.02 (m, 9H). ESI [M+H]^+^: 471.1.

### LYMTAC-3

**6-(6-((2-(2-(2-(3-(4-((((2S,7aR)-7a-(((7-(8-ethyl-7-fluoro-3-hydroxynaphthalen-1-yl)-8-fluoro-4-((R)-3-hydroxy-3-methylpiperidin-1-yl) pyrido [4,3-d] pyrimidin-2-yl) oxy) methyl) hexahydro-1H-pyrrolizin-2-yl) oxy) methyl)-1H-1,2,3-triazol-1-yl) propoxy) ethoxy) ethoxy) ethyl) carbamoyl) pyridin-3-yl)-N-methyl-4-(phenylamino) quinoline-3-carboxamide**

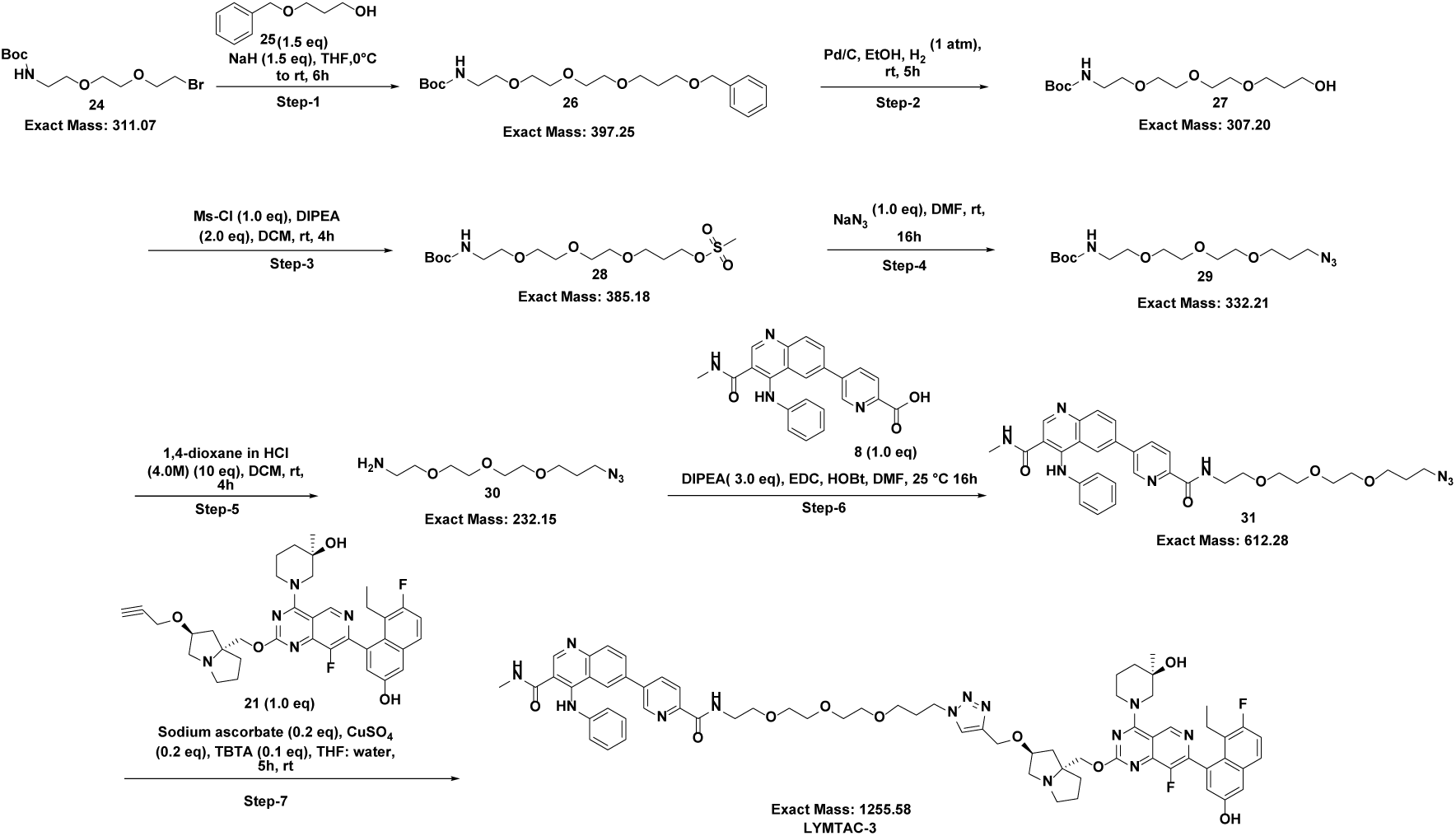

**Step-1: Synthesis of Tert-butyl (1-phenyl-2,6,9,12-tetraoxatetradecan-14-yl) carbamate**

To a 100-mL round-bottomed flask was added 3-(benzyloxy) propan-1-ol (2.396 g, 14.41 mmol, 1.5 eq) in tetrahydrofuran (30 mL) followed by NaH (0.576 g, 14.41 mmol, 1.5 eq) at 0°C and stirred for 1h at same temperature. Then a solution of tert-butyl (2-(2-(2-bromoethoxy) ethoxy) ethyl) carbamate (3.0 g, 9.61 mmol, 1.0 eq) in THF (10 mL) was added to the reaction mixture at 0°C. The reaction mixture was stirred at room temperature for 4h. After completion, the reaction mixture was diluted with cold water (25 mL) and extracted with EtOAc (1 x 50 mL), organic extract was washed with brine (1 x 20 mL) and dried over Na2SO4, filtered and concentrated in vacuo to get the crude material as a colourless oil. The crude material was absorbed onto a plug of silica gel and purified by flash chromatography through a Redi-Sep pre-packed silica gel column (80 g), eluting with a gradient of 15% to 20% EtOAc in hexane, to provide tert-butyl (1-phenyl-2,6,9,12-tetraoxatetradecan-14-yl) carbamate (1.1 g, 2.77 mmol, 28.8 % yield) as a colourless oil. ^1^H NMR (400 MHz, DMSO-*d*6,): δ 7.57 – 7.15 (m, 5H), 6.74 (s, 1H), 4.45 (s, 2H), 3.48 (dddd, *J* = 10.2, 8.9, 4.9, 3.2 Hz, 12H), 3.37 (t, *J* = 6.1 Hz, 2H), 3.06 (q, *J* = 6.0 Hz, 2H), 1.77 (p, *J* = 6.4 Hz, 2H), 1.37 (s, 9H). ESI [M-100]: 298.1.

**Step-2: Synthesis of tert-butyl (2-(2-(2-(3-hydroxypropoxy) ethoxy) ethoxy) ethyl) carbamate**

To a 100-mL round-bottomed flask was added tert-butyl (1-phenyl-2,6,9,12-tetraoxatetradecan-14-yl) carbamate (1.1 g, 2.77 mmol, 1.0 eq) in ethanol (30 mL). After degassing, Pd-C (10%) (0.33 g, 0.310 mmol, 0.11 eq) was added at room temperature and stirred at room temperature for 5 h under hydrogen atmosphere. After completion, the reaction mass was filtered through celite and concentrated to give crude tert-butyl (2-(2-(2-(3-hydroxypropoxy) ethoxy) ethoxy) ethyl) carbamate (0.85 g, 2.77 mmol, 100 % yield) as colorless oil. ^1^H NMR (400 MHz, DMSO-*d*6,): δ 6.75 (d, *J* = 6.4 Hz, 1H), 4.37 (t, *J* = 5.2 Hz, 1H), 3.54 – 3.42 (m, 12H), 3.38 (t, *J* = 6.1 Hz, 2H), 3.06 (q, *J* = 6.0 Hz, 2H), 1.64 (p, *J* = 6.4 Hz, 2H), 1.38 (s, 9H). ESI [M-100]: 208.1

**Step-3: Synthesis of 2,2-dimethyl-4-oxo-3,8,11,14-tetraoxa-5-azaheptadecan-17-yl methane sulfonate**

To a 50-mL round-bottomed flask was added tert-butyl (2-(2-(2-(3-hydroxypropoxy) ethoxy) ethoxy) ethyl) carbamate (250 mg, 0.813 mmol, 1.0 eq), DIPEA (0.3 mL, 1.718 mmol, 2.0 eq) in dichloromethane (30 mL) followed by addition of a solution of MS-Cl (0.063 mL, 0.813 mmol, 1.0 eq) in DCM (1.0 mL) to the reaction mixture at 0°C. The reaction mixture was stirred at room temperature for 6 h. After completion, the reaction mixture was diluted with saturated NaHCO3 solution (20 mL) and extracted with CH2Cl2 (1 x 10 mL). The organic layer was washed with brine (1 x 10 mL) and dried over Na2SO4. The solution was filtered and concentrated in vacuo to give the crude 2,2-dimethyl-4-oxo-3,8,11,14-tetraoxa-5-azaheptadecan-17-yl methane sulfonate (300 mg, 0.778 mmol, 96 % yield) as a colourless oil. ESI [M-100]: 286.2

**Step-4: Synthesis of tert-butyl (2-(2-(2-(3-azidopropoxy) ethoxy) ethoxy) ethyl) carbamate**

To a 25-mL round-bottomed flask was added 2,2-dimethyl-4-oxo-3,8,11,14-tetraoxa-5-azaheptadecan-17-yl methane sulfonate (300 mg, 0.778 mmol) and sodium azide (50.6 mg, 0.778 mmol) in N, N-dimethylformamide (5.0 mL) and stirred at room temperature for 24 h. After completion, the reaction mixture was diluted with cold water (20 mL) and extracted with MTBE (2 x 20 mL). The organic extract was washed with cold water (3 x 10 mL), brine (1 x 10 mL) and dried over Na2SO4. The solution was filtered and concentrated in vacuo at 30°C to give the crude material tert-butyl (2-(2-(2-(3-azidopropoxy) ethoxy) ethoxy) ethyl) carbamate (250 mg, 0.752 mmol, 97 % yield) as a colourless oil, which was used in the next step without any further purification. ESI [M-100]: 233.1

**Step-5: Synthesis of 2-(2-(2-(3-azido propoxy) ethoxy) ethoxy) ethan-1-amine hydrochloride**

To a 50-mL round-bottomed flask was added tert-butyl (2-(2-(2-(3-azidopropoxy) ethoxy) ethoxy) ethyl) carbamate (250 mg, 0.752 mmol) in dichloromethane (20 mL) followed by addition of 4 M HCl in 1,4-dioxane (1880 µL, 7.52 mmol, 10 eq) at 0°C. The reaction mixture was stirred at room temperature for 4 h. After completion, the reaction mixture was concentrated and the crude 2-(2-(2-(3-azido propoxy) ethoxy) ethoxy) ethan-1-amine hydrochloride (190 mg) was used in the next step without any further purification. ESI [M+H]^+^: 233.1

**Step-6: Synthesis of 6-(6-((2-(2-(2-(3-azido propoxy) ethoxy) ethoxy) ethyl) carbamoyl) pyridin-3-yl)-N-methyl-4-(phenylamino) quinoline-3-carboxamide**

To a 50-mL round-bottomed flask was added 2-(2-(2-(3-azido propoxy) ethoxy) ethoxy) ethan-1-amine hydrochloride (186 mg, 0.693 mmol, 1.2 eq), 5-(3-(methylcarbamoyl)-4-(phenylamino) quinolin-6-yl) picolinic acid (230 mg, 0.577 mmol, 1.0 eq) and DIPEA (605 µL, 3.46 mmol, 6.0 eq) in N, N-dimethylformamide (10 mL), followed by addition of EDC (166 mg, 0.866 mmol, 1.5 eq) and 1H-benzo[d] [1,2,3] triazol-1-ol (117 mg, 0.866 mmol, 1.5 eq) to the reaction mixture at 0°C. The reaction mixture was stirred at room temperature for 16 h. After completion, the reaction mixture was diluted with cold water (30 mL) and extracted with EtOAc (2 x 20 mL). The organic extract was washed with cold water (3 x 20 mL), brine solution (1 x 10 mL) and dried over Na2SO4. The solution was filtered and concentrated in vacuo to provide crude 6-(6-((2-(2-(2-(3-azido propoxy) ethoxy) ethoxy) ethyl) carbamoyl) pyridin-3-yl)-N-methyl-4-(phenylamino) quinoline-3-carboxamide (300 mg) as a light-yellow solid, which was used next step without further purification. ESI [M+H]^+^: 613.2

**Step-7: Synthesis of 6-(6-((2-(2-(2-(3-(4-((((2S,7aR)-7a-(((7-(8-ethyl-7-fluoro-3-hydroxynaphthalen-1-yl)-8-fluoro-4-((R)-3-hydroxy-3-methylpiperidin-1-yl) pyrido [4,3-d] pyrimidin-2-yl) oxy) methyl) hexahydro-1H-pyrrolizin-2-yl) oxy) methyl)-1H-1,2,3-triazol-1-yl) propoxy) ethoxy) ethoxy) ethyl) carbamoyl) pyridin-3-yl)-N-methyl-4-(phenylamino) quinoline-3-carboxamide (LYMTAC-3)**

To a 10-mL round-bottomed flask was added 6-(6-((2-(2-(2-(3-azidopropoxy)ethoxy)ethoxy)ethyl)carbamoyl)pyridin-3-yl)-N-methyl-4-(phenylamino)quinoline-3-carboxamide (25 mg, 0.041 mmol, 1.0 eq) and (R)-1-(7-(8-ethyl-7-fluoro-3-hydroxynaphthalen-1-yl)-8-fluoro-2-(((2S,7aR)-2-(prop-2-yn-1-yloxy)tetrahydro-1H-pyrrolizin-7a(5H)-yl)methoxy)pyrido[4,3-d]pyrimidin-4-yl)-3-methylpiperidin-3-ol (26.3 mg, 0.041 mmol, 1.0 eq) in tetrahydrofuran (4.0 mL) and water (1.0 mL), followed by addition of copper(II) sulfate (2.60 mg, 0.016 mmol, 0.4 eq) and TBTA (4.33 mg, 8.16 µmol, 0.2 eq), (+)-sodium L-ascorbate (3.23 mg, 0.016 mmol, 0.4 eq) to the reaction mixture at 0°C. The reaction mixture was stirred at room temperature for 5 h. After completion, the reaction mixture was purified by Prep-HPLC, using column Kinetex EVO C18 (250×21.2) mm 5.0µm, mobile phase A: 0.1% Formic acid in water B: ACN, flow rate 15ml/ min), to give 6-(6-((2-(2-(2-(3-(4-((((2S,7aR)-7a-(((7-(8-ethyl-7-fluoro-3-hydroxynaphthalen-1-yl)-8-fluoro-4-((R)-3-hydroxy-3-methylpiperidin-1-yl)pyrido[4,3-d]pyrimidin-2-yl)oxy)methyl)hexahydro-1H-pyrrolizin-2-yl)oxy)methyl)-1H-1,2,3-triazol-1-yl)propoxy)ethoxy)ethoxy)ethyl)carbamoyl)pyridin-3-yl)-N-methyl-4-(phenylamino)quinoline-3-carboxamide (23 mg, 0.018 mmol, 44.9 % yield) as a pale yellow solid. ^1^H NMR (400 MHz, DMSO-*d*6, 400 MHz): δ 10.06 (s, 1H), 9.97 (s, 1H), 9.22 (s, 1H), 8.82 (s, 1H), 8.77 (s, 1H), 8.72 (t, *J* = 5.8 Hz, 1H), 8.52 (s, 1H), 8.37 (s, 1H), 8.17 (dd, *J* = 17.0, 8.7 Hz, 2H), 8.10 – 8.01 (m, 3H), 7.81 – 7.73 (m, 1H), 7.40 – 7.26 (m, 4H), 7.13 – 7.01 (m, 4H), 4.76 (d, *J* = 7.0 Hz, 1H), 4.47 (s, 2H), 4.36 (t, *J* = 7.0 Hz, 3H), 4.18 (s, 1H), 4.10 – 3.95 (m, 3H), 3.67 – 3.41 (m, 16H), 3.06 (d, *J* = 4.8 Hz, 1H), 2.94 (s, 1H), 2.73 (d, *J* = 14.6 Hz, 2H), 2.56 (d, *J* = 4.3 Hz, 3H), 2.15 (dd, *J* = 13.2, 6.0 Hz, 2H), 2.06 – 1.94 (m, 3H), 1.86 (s, 3H), 1.79 – 1.58 (m, 6H), 1.16 (d, *J* = 9.2 Hz, 3H), 0.73 (q, *J* = 7.4 Hz, 3H). ESI [M+H]^+^: 1256.2

### LYMTAC-4

**6-(6-(3-(4-((((2S,7aR)-7a-(((7-(8-ethyl-7-fluoro-3-hydroxynaphthalen-1-yl)-8-fluoro-4-((R)-3-hydroxy-3-methylpiperidin-1-yl)pyrido[4,3-d] pyrimidin-2-yl) oxy) methyl) hexahydro-1H-pyrrolizin-2-yl) oxy) methyl)-1H-1,2,3-triazol-1-yl) azetidine-1-carbonyl) pyridin-3-yl)-N-methyl-4-(phenylamino) quinoline-3-carboxamide**

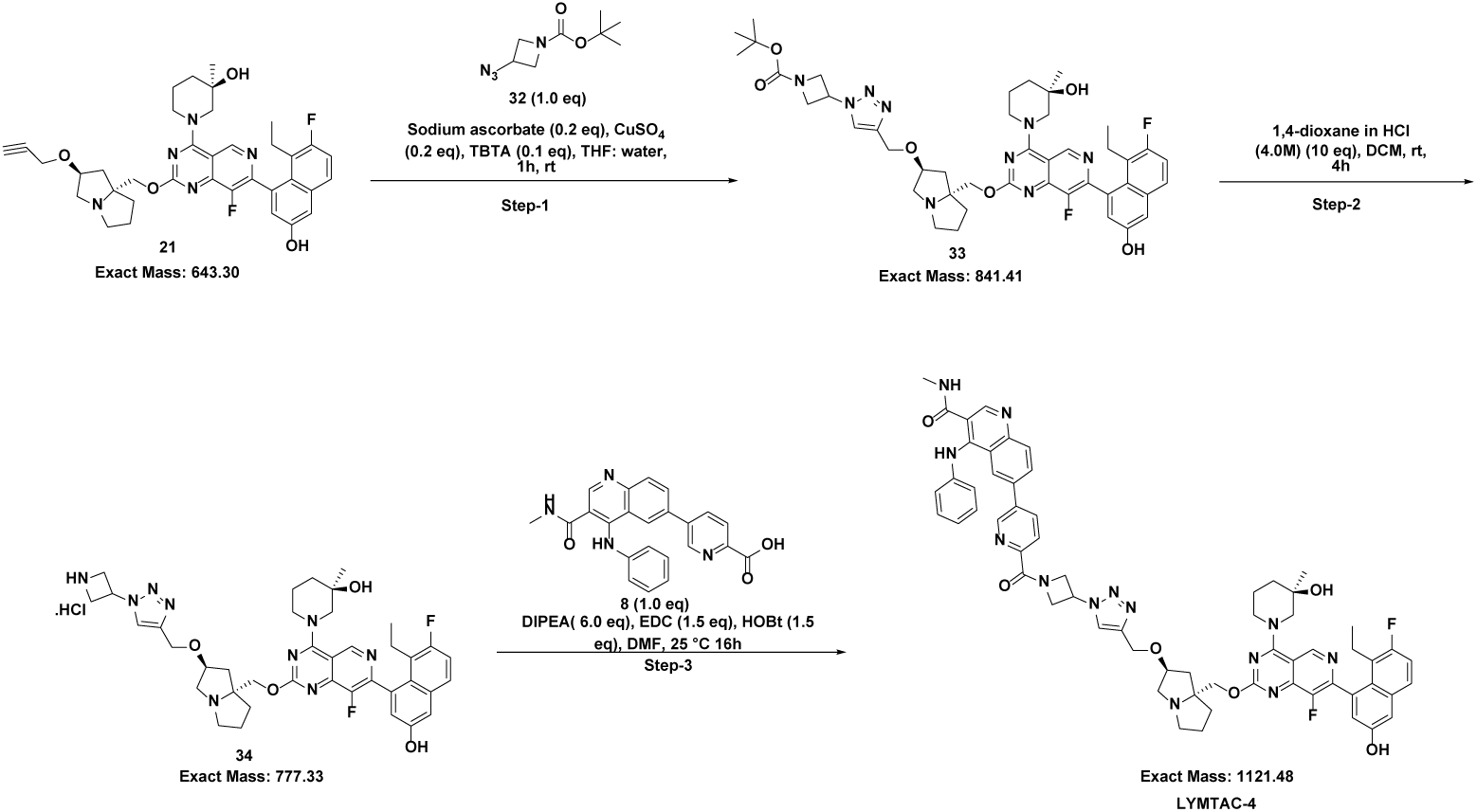

**Step-1: Synthesis of tert-butyl 3-(4-((((2S,7aR)-7a-(((7-(8-ethyl-7-fluoro-3-hydroxynaphthalen-1-yl)-8-fluoro-4-((R)-3-hydroxy-3-methylpiperidin-1-yl) pyrido[4,3-d] pyrimidin-2-yl) oxy) methyl) hexahydro-1H-pyrrolizin-2-yl) oxy) methyl)-1H-1,2,3-triazol-1-yl) azetidine-1-carboxylate**

To a 10-mL round-bottomed flask was added (R)-1-(7-(8-ethyl-7-fluoro-3-hydroxynaphthalen-1-yl)-8-fluoro-2-(((2S,7aR)-2-(prop-2-yn-1-yloxy)tetrahydro-1H-pyrrolizin-7a(5H)-yl)methoxy)pyrido[4,3-d]pyrimidin-4-yl)-3-methylpiperidin-3-ol (50 mg, 0.078 mmol, 1.0 eq, synthesis described in KRAS PROTAC) and tert-butyl 3-azidoazetidine-1-carboxylate (15.40 mg, 0.078 mmol, 1.0 eq) in tetrahydrofuran (4.0 mL) and water (1.0 mL), followed by addition of copper(II) sulfate (4.96 mg, 0.031 mmol, 0.2 eq) and TBTA (8.23 mg, 0.016 mmol, 0.1 eq), (+)-sodium L-ascorbate (6.15 mg, 0.031 mmol, 0.2 eq) to the reaction mixture at 0°C. The reaction mixture was stirred at room temperature for 1 h. After completion, the reaction mixture was filtered through celite and washed with ethyl acetate (20 mL), the organic extract was washed with brine (1x 5 mL) and dried over Na2SO4. The solution was filtered and concentrated in vacuo to give the crude of tert-butyl 3-(4-((((2S,7aR)-7a-(((7-(8-ethyl-7-fluoro-3-hydroxynaphthalen-1-yl)-8-fluoro-4-((R)-3-hydroxy-3-methylpiperidin-1-yl) pyrido [4,3-d] pyrimidin-2-yl) oxy) methyl) hexahydro-1H-pyrrolizin-2-yl) oxy) methyl)-1H-1,2,3-triazol-1-yl) azetidine-1-carboxylate (70 mg) as a light-yellow oil, which was used for next step without further purification. ESI [M+H]+: 842.3

**Step-2: Synthesis of (R)-1-(2-(((2S,7aR)-2-((1-(azetidin-3-yl)-1H-1,2,3-triazol-4-yl) methoxy) tetrahydro-1H-pyrrolizin-7a(5H)-yl) methoxy)-7-(8-ethyl-7-fluoro-3-hydroxynaphthalen-1-yl)-8-fluoropyrido[4,3-d] pyrimidin-4-yl)-3-methylpiperidin-3-ol hydrochloride**

To a 25-mL round-bottomed flask was added tert-butyl 3-(4-((((2S,7aR)-7a-(((7-(8-ethyl-7-fluoro-3-hydroxynaphthalen-1-yl)-8-fluoro-4-((R)-3-hydroxy-3-methylpiperidin-1-yl) pyrido [4,3-d] pyrimidin-2-yl) oxy) methyl) hexahydro-1H-pyrrolizin-2-yl) oxy) methyl)-1H-1,2,3-triazol-1-yl) azetidine-1-carboxylate (65 mg, 0.077 mmol, 1.0 eq) in dichloromethane (10 mL) followed by addition of 4 M HCl in 1,4 dioxane (193 µL, 0.772 mmol, 10.0 eq) to the reaction mixture at 0°C and stirred at room temperature for 4 h. After completion, the reaction mass was concentrated and the crude (R)-1-(2-(((2S,7aR)-2-((1-(azetidin-3-yl)-1H-1,2,3-triazol-4-yl) methoxy) tetrahydro-1H-pyrrolizin-7a(5H)-yl) methoxy)-7-(8-ethyl-7-fluoro-3-hydroxynaphthalen-1-yl)-8-fluoropyrido[4,3-d] pyrimidin-4-yl)-3-methylpiperidin-3-ol hydrochloride compound (70 mg) was used directly in the next reaction without purification. ESI [M+H]+: 742.1

**Step-3: Synthesis of 6-(6-(3-(4-((((2S,7aR)-7a-(((7-(8-ethyl-7-fluoro-3-hydroxynaphthalen-1-yl)-8-fluoro-4-((R)-3-hydroxy-3-methylpiperidin-1-yl) pyrido[4,3-d] pyrimidin-2-yl) oxy) methyl) hexahydro-1H-pyrrolizin-2-yl) oxy) methyl)-1H-1,2,3-triazol-1-yl) azetidine-1-carbonyl) pyridin-3-yl)-N-methyl-4-(phenylamino) quinoline-3-carboxamide (LYMTAC-4)**

To a 10-mL round-bottomed flask was added 5-(3-(methylcarbamoyl)-4-(phenylamino)quinolin-6-yl)picolinic acid (25 mg, 0.063 mmol, 1.0 eq) and (R)-1-(2-(((2S,7aR)-2-((1-(azetidin-3-yl)-1H-1,2,3-triazol-4-yl)methoxy)tetrahydro-1H-pyrrolizin-7a(5H)-yl)methoxy)-7-(8-ethyl-7-fluoro-3-hydroxynaphthalen-1-yl)-8-fluoropyrido[4,3-d]pyrimidin-4-yl)-3-methylpiperidin-3-ol hydrochloride (48.8 mg, 0.063 mmol, 1.0 eq), DIPEA (65.8 µL, 0.376 mmol, 6.0 eq) in N, N-dimethylformamide (5.0 mL) followed by addition of EDC (18.04 mg, 0.094 mmol, 1.5 eq) and 1H-benzo[d][1,2,3]triazol-1-ol (12.72 mg, 0.094 mmol, 1.5 eq) to the reaction mixture at 0°C and stirred at room temperature for 16 h. After completion, the reaction mixture was concentrated and purified by Prep-HPLC purification (Column X-Select C18 (250×19) mm 5.0µm, mobile phase A: 10 Mm Ammonium acetate in water B: ACN, flow rate 15ml/min), to provide 6-(6-(3-(4-((((2S,7aR)-7a-(((7-(8-ethyl-7-fluoro-3-hydroxynaphthalen-1-yl)-8-fluoro-4-((R)-3-hydroxy-3-methylpiperidin-1-yl)pyrido[4,3-d] pyrimidin-2-yl) oxy) methyl) hexahydro-1H-pyrrolizin-2-yl) oxy) methyl)-1H-1,2,3-triazol-1-yl) azetidine-1-carbonyl) pyridin-3-yl)-N-methyl-4-(phenylamino) quinoline-3-carboxamide (14 mg, 0.012 mmol, 19.88 % yield) as a yellow solid. ^1^H NMR (400 MHz, DMSO-*d*6): δ 9.94 (s, 2H), 9.22 (d, *J*=1.4 Hz, 1H), 8.89 (s, 1H), 8.78 (s, 1H), 8.51 – 8.38 (m, 3H), 8.27 – 8.14 (m, 2H), 8.07 (dd, *J*=15.1, 8.5 Hz, 2H), 7.76 (dd, *J*=9.1, 6.0 Hz, 1H), 7.41 – 7.23 (m, 4H), 7.14 – 6.97 (m, 4H), 5.62 (p, *J*=7.3, 6.7 Hz, 1H), 5.20 (t, *J*=9.4 Hz, 1H), 4.95 (dd, *J*=11.0, 5.3 Hz, 1H), 4.75 (d, *J*=6.7 Hz, 1H), 4.71 – 4.62 (m, 1H), 4.53 (s, 2H), 4.46 – 4.26 (m, 2H), 4.24 (t, *J*=5.2 Hz, 1H), 4.14 – 3.95 (m, 3H), 3.58 (dd, *J*=41.6, 13.3 Hz, 2H), 3.11 (dd, *J*=10.9, 4.7 Hz, 1H), 2.98 (dd, *J*=9.8, 5.3 Hz, 1H), 2.76 (dq, *J*=15.3, 6.1, 4.7 Hz, 2H), 2.16 – 2.15 (m, 1H) 2.18 (dd, *J*=13.3, 5.9 Hz, 3H), 1.86 (s, 4H), 1.83 – 1.61 (m, 7H), 1.17 (d, *J*=8.7 Hz, 3H), 0.73 (q, *J*=7.4 Hz, 3H). ESI [M+H]+: 1122.3

